# Neural trajectories in the supplementary motor area and primary motor cortex exhibit distinct geometries, compatible with different classes of computation

**DOI:** 10.1101/650002

**Authors:** Abigail A. Russo, Ramin Khajeh, Sean R. Bittner, Sean M. Perkins, John P. Cunningham, Laurence F. Abbott, Mark M. Churchland

## Abstract

The supplementary motor area (SMA) is believed to contribute to higher-order aspects of motor control. To examine this contribution, we employed a novel cycling task and leveraged an emerging strategy: testing whether population trajectories possess properties necessary for a hypothesized class of computations. We found that, at the single-neuron level, SMA exhibited multiple response features absent in M1. We hypothesized that these diverse features might contribute, at the population level, to avoidance of ‘population trajectory divergence’ – ensuring that two trajectories never followed the same path before separating. Trajectory divergence was indeed avoided in SMA but not in M1. Network simulations confirmed that low trajectory divergence is necessary when guidance of future action depends upon internally tracking contextual factors. Furthermore, the empirical trajectory geometry – helical in SMA versus elliptical in M1 – was naturally reproduced by networks that did, versus did not, internally track context.

## Introduction

The supplementary motor area (SMA) is implicated in higher-order aspects of motor control^1–3^. SMA lesions cause motor neglect^4,5^, unintended utilization^6^, and difficulty performing temporal sequences^7–9^. Relative to primary motor cortex (M1), SMA activity is less coupled to actions of a specific body part^10–12^. Instead, SMA computations appear related to learned sensory-motor associations^11^, reward anticipation^13^, internal initiation and guidance of movement^3,14^, movement timing^15,16^, and movement sequencing^7,17,18^. Single-neuron responses in SMA reflect a variety of task-specific contingencies. For example, in a sequence of three movements, a neuron may burst only when pulling precedes pushing. Another neuron might respond before the third movement regardless of the particular sequence^19^. Different response features are observed in different tasks. Single SMA neurons exhibit a mixture of ramping and rhythmic activity during an interval timing task^20^, and the SMA population exhibits amplitude-modulated circular trajectories during rhythmic tapping^21^. A common thread linking prior studies is that SMA computations are hypothesized to be critical when pending action depends upon internal, abstract, and/or contextual factors. An important challenge is linking these high-level ideas to network-level implementations. What general properties should activity exhibit in networks performing the hypothesized class of computations?

There exist many quantitative methods for relating population activity and computation (e.g.,^22–24^). These include decoding key hypothesized signals (e.g., via regression^25^), or directly comparing empirical and simulated population activity (e.g., via canonical correlation^26^). An emerging strategy is to consider the geometry of the population response: the arrangement of population states across conditions^23,27,28^ and/or the shape traced by activity in neural state-space^15,26,29–36^. A given geometry may be consistent with some types of computation but not others^37^. An advantage of this approach is that it is sometimes possible to measure geometric properties that are expected to hold for a class of computations, regardless of the exact instantiation. For example, we recently characterized M1 activity using a metric, ‘trajectory tangling’, that assesses whether activity could be generated by noise-robust network dynamics^38^. This approach revealed a population-level property that was conserved across tasks and species.

Here we consider the hypothesis that SMA guides movement by internally tracking contextual factors, and derive a prediction regarding the population trajectory geometry appropriate for that class of computations. We predict that SMA trajectories should avoid ‘divergence’; trajectories should be structured, across time and across conditions, such that it is never the case that two trajectories follow the same path and then separate. Low divergence is essential to ensure that neural activity can distinguish between situations with different future motor outputs, even if the present motor output is similar. We tested whether this hypothesized geometry is indeed observed in a novel task, and whether it can account for single-neuron response properties and task-specific population-level features.

We employed a recently developed cycling task that shares some features with sequence / timing tasks but involves continuous motor output and thus provides a novel perspective on SMA response properties. We found that the population response in SMA, but not M1, exhibited low trajectory divergence. The major features of SMA responses, both at the population and single-neuron levels, could be understood as serving to maintain low divergence. Simulations confirmed that low divergence was necessary for a network to guide action based on internal / contextual information. Furthermore, artificial networks naturally adopted low-divergence, SMA-like trajectories when performing computations that required internally tracking contextual factors. Thus, a broad hypothesis regarding the type of computation performed by SMA successfully predicts SMA population trajectory geometry, and provides an explanation for seemingly diverse task-specific response features.

## Results

### Task and behavior

We trained two rhesus macaque monkeys to grasp a hand-pedal and cycle through a virtual landscape^38^ (Fig. 1a). Each trial required the monkey to cycle between a pair of targets. The trial began with the virtual position stationary on the first target, with the pedal orientation either straight up (‘top-start’) or straight down (‘bottom-start’). After a 1000 ms hold period, the second target appeared. Second-target distance determined the number of revolutions that had to be performed: 1, 2, 4, or 7 cycles. Following a 500-1000 ms randomized delay period, a go-cue (brightening of the second target) was delivered. The monkey then cycled to that target and remained stationary to receive a juice reward. Because targets were separated by an integer number of cycles, the second target was acquired with the same pedal orientation (straight up or down) as for the first target. Landscape color indicated whether forward virtual motion required ‘forward’ cycling (the hand moved away from the body at the top of the cycle) or ‘backward’ cycling (the hand moved toward the body at the top of the cycle). Using a block-randomized design, monkeys performed all combinations of two cycling directions, two starting orientations, and four cycling distances. Averages of hand kinematics, muscle activity and neural activity were computed after temporal alignment to account for small trial-by-trial differences in cycling speed^38^.

**Figure 1.**
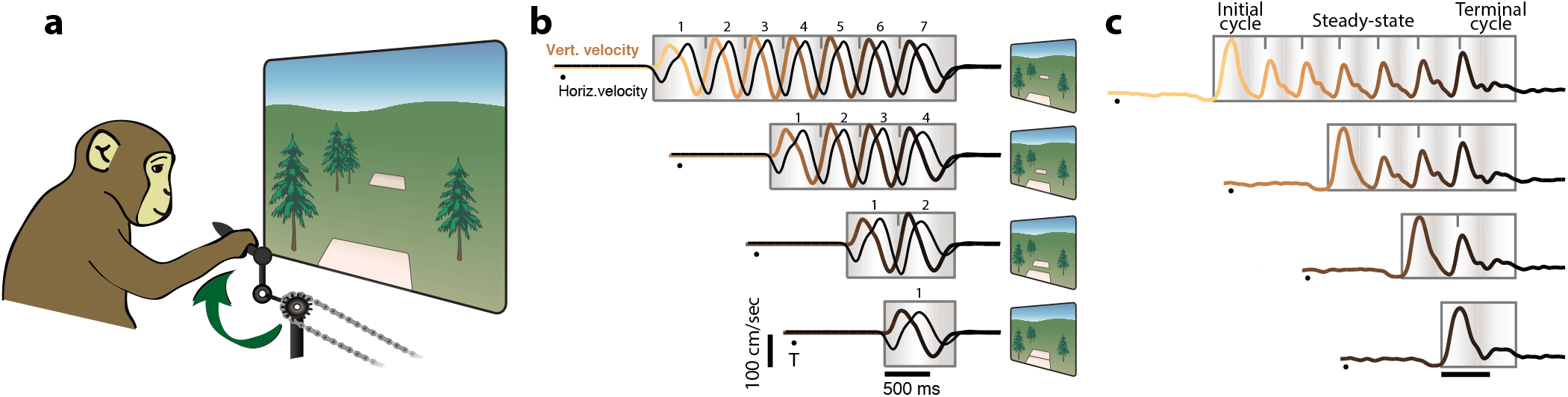
Task schematic, behavior, and muscle activity during cycling. **a)** Schematic of the cycling task. Monkeys grasped a hand pedal and cycled through a virtual environment for a number of cycles prescribed by target distance. The schematic illustrates forward cycling, instructed by a green environment. Backward cycling was instructed by an orange, desert-like environment. **b)** Trial-averaged hand velocity (vertical and horizontal) for seven-cycle, four-cycle, two-cycle and one-cycle movements. Data are for forward cycling, starting at cycle’s bottom (monkey C). Vertical velocity traces are colored from tan to black to indicate time with respect to the end of movement. Black dots indicate the time of target appearance. Gray box with shading indicates the period of time where the pedal was moving (preceding go cue is not shown). Shading indicates vertical hand position; light shading indicates cycle apex. Tick marks indicate cycle divisions (used for analysis, and not an overt aspect of the task or behavior). Task schematic panels (right) indicate how target distance indicated the number of cycles to be produced. **c)** Muscle activity, recorded from the medial head of the triceps (monkey D). Intra-muscularly recorded voltages were rectified, filtered, and trial averaged. Data are shown for the four distances, backward cycling, starting at cycle’s top. Same plotting conventions as in b. Labels at top indicate, for the seven-cycle movement, the initial cycle, the terminal cycle, and the intervening steady-state cycles.

Vertical and horizontal hand velocity displayed nearly sinusoidal temporal profiles (Fig. 1b). Muscle activity patterns (Fig. 1c) were often non-sinusoidal, and initial-cycle and/or terminal-cycle patterns often departed from the middle-cycle pattern (e.g., the initial-cycle response is larger for the example shown). This is an expected consequence of the need to accelerate the arm when starting and to decelerate the arm when stopping. Muscle activity and hand kinematics differed in many ways, yet shared the following property: the response when cycling a given distance was a concatenation of an initial-cycle response, some number of middle cycles with a repeating response, and a terminal-cycle response. We refer to the middle cycles as ‘steady-state’ cycling, reflecting the fact that kinematics and muscle activity repeated across such cycles, both within a cycling distance and across distances. Seven-cycle movements had ~5 steady-state cycles and four-cycle movements had ~2 steady-state cycles. Two- and one-cycle movements involved little or no steady-state cycling. Such structure is reminiscent of a sequence task (e.g., a four-cycle movement follows an ABBC pattern). However, both movement and accompanying muscle activity were continuous; cycle divisions are employed simply for presentation and analysis.

Our motivating hypothesis, derived from prior studies^7,11,14–16,19–21,39–41^, is that SMA tracks internal and/or contextual factors for the purpose of guiding action. If so, the SMA population response should be shaped by the need to consistently distinguish situations that involve different future actions, even if the current motor output is identical. The cycling task produced multiple instances of this scenario, both within and between conditions. Consider the second and fifth cycles of a seven-cycle movement (Fig. 1b,c). Motor output is essentially identical at these two phases of the task. Yet in two more cycles the output will differ. The same is true when comparing the second cycle of seven-cycle and four-cycle movements. Does the need to distinguish between such situations account for the geometry of the SMA population response? While this is fundamentally a population-level question, we begin by examining single-neuron responses. We then describe specific features of the population response. Finally, we consider a general property of population trajectory geometry required when a network must internally keep track of context. We use simulations to validate that approach, and then test whether the key predictions hold for the empirical trajectories.

### Single-neuron responses

Well-isolated single neurons were recorded sequentially from SMA (77 and 70 recordings for monkeys C and D) and M1 (109 and 103 recordings). Recording locations were guided via MRI landmarks, microstimulation, light touch, and muscle palpation to confirm the trademark properties of each region. M1 recordings included not only sulcal and surface primary motor cortex (M1 proper) but also recordings from the immediately adjacent aspect of dorsal premotor cortex^38^. Neurons in both SMA and M1 were robustly modulated during cycling. Firing rate modulations (maximum minus minimum rate) averaged 52 and 57 spikes/s for SMA (monkey C and D) and 73 and 64 spikes/s for M1.

In M1, single-neuron responses (Fig 2a-c) were typically complex, yet showed two consistent features. First, for a given cycling distance, responses repeated across steady-state cycles. For example, for a seven-cycle movement, the firing rate profile was very similar across cycles 2-6^38^. Second, response elements – initial-cycle, steady-state, and terminal-cycle responses – were conserved across cycling distances. Thus, although M1 responses rarely matched patterns of muscle activity or kinematics, they shared the same general structure. Responses were essentially a concatenation of an initial-cycle response, a steady-state response, and a terminal-cycle response. Even complex responses that might be mistaken as ‘noise’ displayed this structure (Fig. 2c).

**Figure 2.**
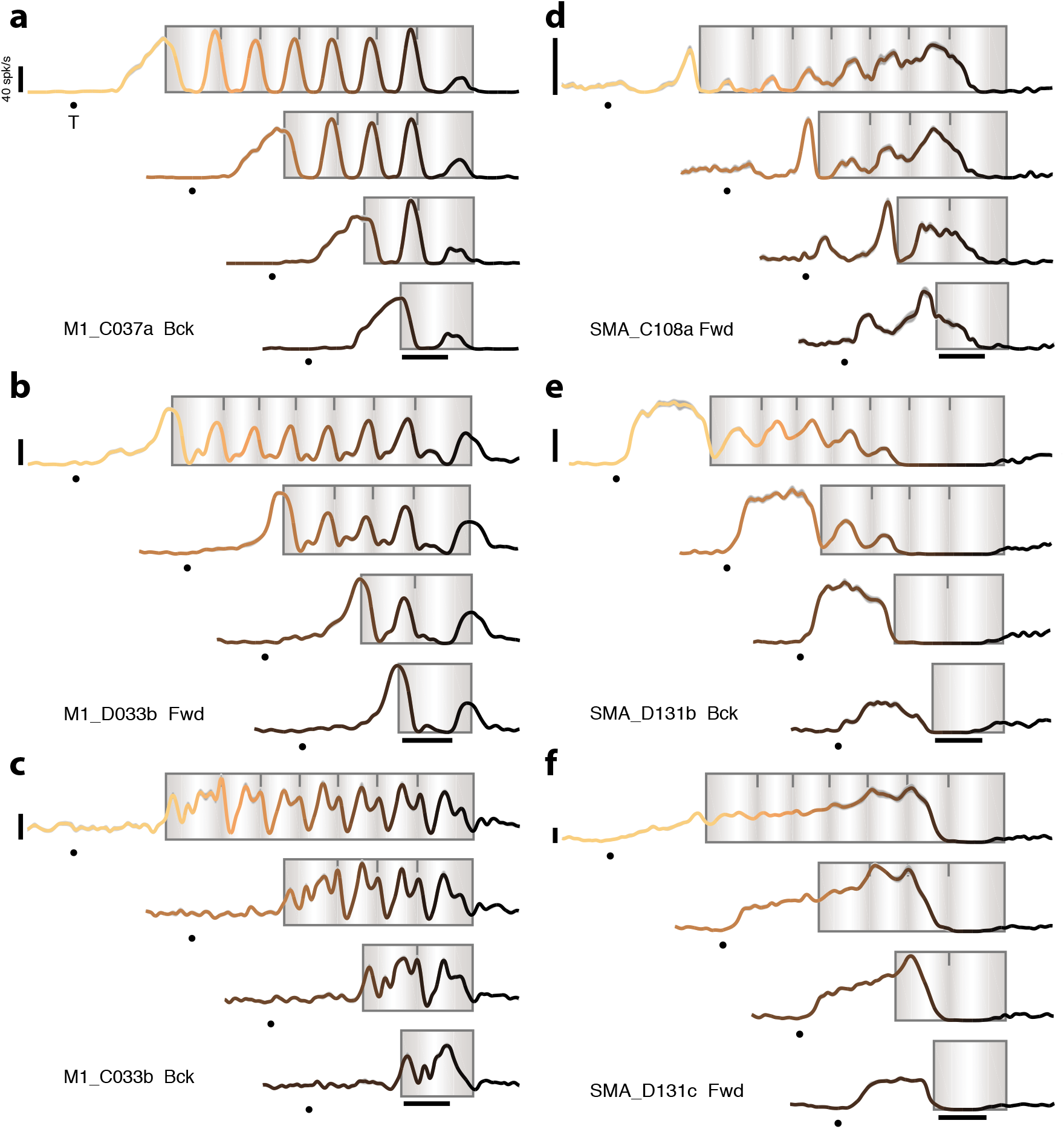
Responses of example neurons. **a-c)** Firing rates for three example M1 neurons. Same plotting conventions as in Fig. 1c. The label in each panel indicates the region, monkey (C or D), and the cycling direction for which the data were recorded. All data are for conditions where cycling started from the bottom position. All calibrations are 40 spikes/s. Gray envelopes around each trace (typically barely visible) give the standard error of the mean. **d-f)** Firing rates for three SMA neurons. Same format as (a-c).

Neurons in SMA (Fig. 2d-f) displayed a different set of properties. Responses were typically a mixture of rhythmic and ramp-like features (Fig. 2d). As a result, during steady-state cycling, single-neuron responses in SMA had a greater proportion of their power well below the ~2 Hz cycling frequency (Fig. 3a,b). Due in part to these slow changes in firing rate, a clear ‘steady-state’ response was rarely reached. Furthermore, the initial-cycle response in SMA often differed across cycling distances (e.g., compare the seven-cycle and two-cycle response in Fig. 2e) even when muscle and M1 responses were similar. Yet terminal-cycle responses were largely preserved across distances. For example, in Figure 2e, the response during a four-cycle movement is similar to that during the last four cycles of a seven-cycle movement.

**Figure 3.**
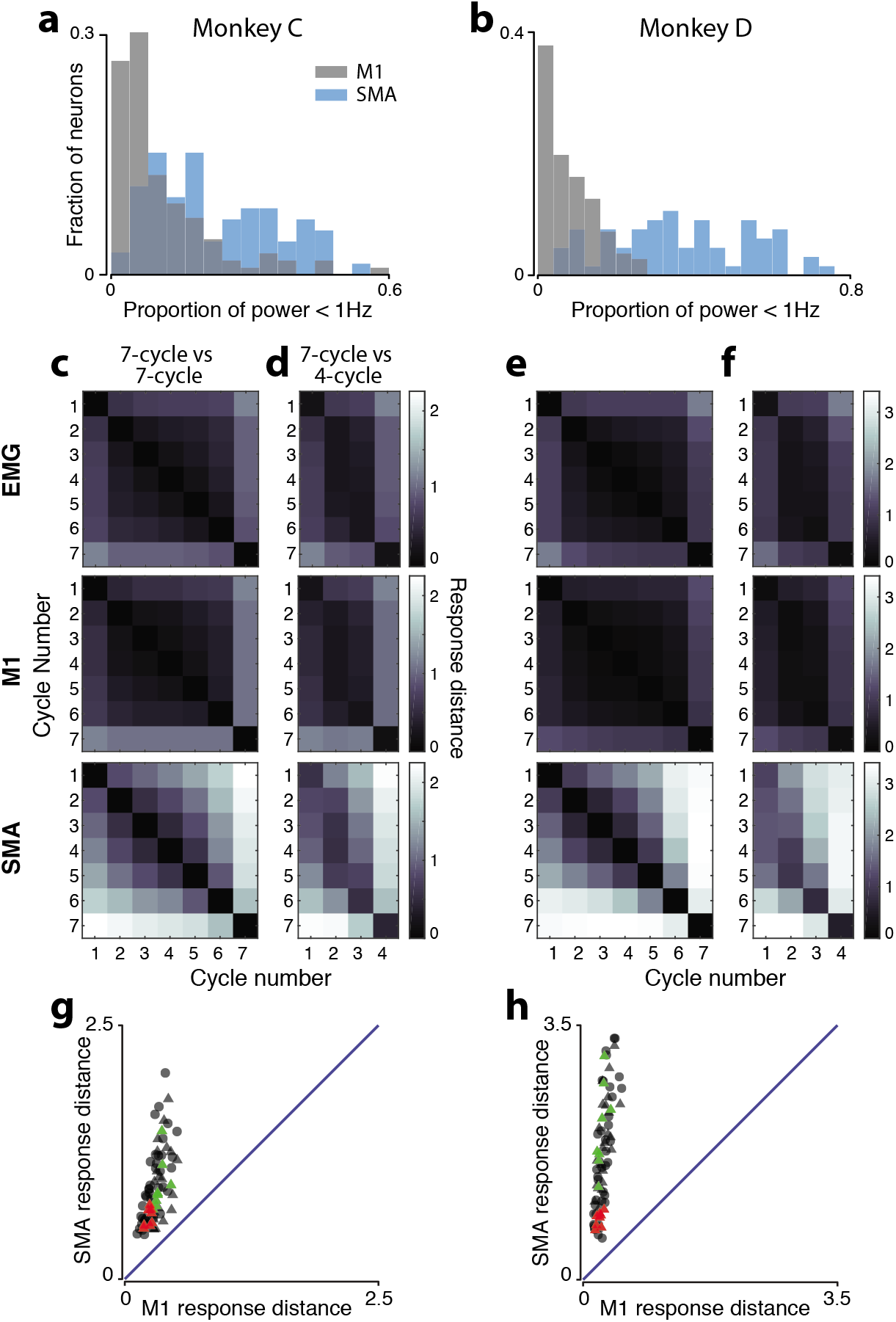
SMA responses show greater cycle-to-cycle differences. **a)** Histogram of the proportion of firing-rate power <1 Hz during steady state cycling. Because cycling occurred at ~2 Hz, power below 1 Hz corresponds to slower across-cycle changes such as firing-rate ramps. For each neuron, power was computed for each of the 7-cycle conditions (two cycling directions by two starting positions). Power was computed after mean centering (ensuring no power at 0 Hz). The proportion of power < 1 Hz was then averaged across conditions to yield one value per neuron. Data are for monkey C. **b)** Same for monkey D. **c)** Matrices of response distances when comparing cycles within a seven-cycle movement. For each comparison (e.g., cycle two versus cycle three) normalized response distance was computed for each of the four conditions (forward and backward cycling, starting at the top and bottom) and then averaged across conditions. Data are for monkey C. Note that the matrix is symmetric; response distance between cycle two and three is the same as between three and two. The diagonal is necessarily zero; the response on cycle three cannot differ from itself when compared within a seven-cycle movement. **d)** Matrices of response distances when comparing seven-cycle and four-cycle movements. Data are for monkey C. Matrices are not symmetric (they are not even square) and there is no diagonal of values that are necessarily zero. **e)** Same as panel c, but for monkey D. **f)** Same as panel d, but for monkey D. **g)** Response distance for SMA versus M1 for all comparisons among steady-state cycles. This includes comparisons within seven-cycle movements (circles) and between seven-cycle and four-cycle movements (triangles). Each symbol corresponds to one comparison; e.g., the third cycle of a four-cycle movement with the fourth cycle of a seven-cycle movement. Data for each condition (different cycling directions and starting positions) are plotted separately. For each of the four conditions, there are ten total comparisons within the seven-cycle movement, and an additional ten when comparing seven- and four-cycle movements, resulting in eighty total comparisons. Red triangles highlight comparisons between cycles equidistant from movement end: six-of-seven versus three-of-four and five-of-seven versus two-of-four. Green triangles highlight comparisons between cycles equidistant from movement beginning: two-of-seven versus two-of-four and three-of-seven versus three-of-four. Data are for monkey C. **h)** Same for monkey D.

### Individual-cycle responses are more distinct in SMA

We compared the response on each cycle with that on every other cycle, both within seven-cycle movements (Fig. 3c,e), and between seven-cycle and four-cycle movements (Fig 3d,f). For each comparison, we defined ‘response distance’ as the root-mean-squared difference in firing rate across all neurons and times within that cycle. Response distance was normalized by the typical intra-cycle firing-rate modulation for that condition. This analysis thus assesses the degree to which responses differ across cycles, relative to the response magnitude within a single cycle. Individual-neuron responses were normalized to avoid analysis being dominated by a few high firing-firing rate neurons. To avoid response distance being inflated by sampling error, we used principal component analysis (PCA) to de-noise the response of each neuron (Methods). Results were not sensitive to the choice of dimensionality so long as it was sufficient to capture a majority of the data variance. Response distance was averaged across the two cycling directions and starting positions (Fig. 3c-f), or shown for each independently (Fig. 3g,h).

Figure 3c-f plots response distance for every comparison in matrix form. For M1 (middle row), responses were similar among all steady-state cycles, resulting in a central dark block. This block is square for the within-seven-cycle comparison and rectangular for the seven-versus-four-cycle comparison. Outer rows and columns are lighter reflecting the fact that initial- and terminal-cycle responses differed both from one another and from steady-state responses. This analysis confirms that M1 responses involve a distinct initial-cycle response, a repeating steady-state response, and a distinct terminal-cycle response. Essentially identical structure was observed for the muscle populations (Fig. 3c-f, top row). These results agree with the finding that M1 activity relates to the execution of the present movement^42–44^.

For SMA, the central block of high similarity was largely absent (Fig. 3c-f, bottom row). Instead, distance grew steadily with temporal separation. For example, within a seven-cycle movement, the second-cycle response was modestly different from the third-cycle response, fairly different from the fifth-cycle response, and very different from the seventh-cycle response. The average response distance between steady-state cycles was 3.1 times larger (monkey C) and 6.1 times larger (monkey D) for SMA versus M1 (p<0.0001 via bootstrap for each monkey). SMA showed dissimilar responses across steady-state cycles both within a cycling distance (Fig. 3c,e), and when comparing across cycling distances (Fig. 3d,f) (p<0.0001 in all cases).

For all comparisons among steady-state-cycles, across all conditions, response distance was higher for SMA (Fig. 3g,h, p<10^−10^ via paired t-test for each monkey). Yet the magnitude of this effect depended on the situation. Intriguingly, SMA response distance was modest when comparing cycles equidistant from movement end (red triangles) – e.g., cycle six-of-seven versus three-of-four. SMA response distance was not similarly low when comparing cycles equidistant from movement beginning (green triangles) – e.g., cycle two-of-seven versus two-of-four. This same effect can be observed in Figure 3d,f (bottom): the diagonal ending in the bottom-right corner contains smaller values (darker squares) than the diagonal beginning in the top-left corner. As a result, response distance in SMA was significantly smaller when comparing the last three cycles versus the first three cycles (p<0.001, for each monkey, bootstrap). This asymmetry was greater for SMA (p<0.05 for monkey C and p<0.0001 for monkey D) than for M1, where responses distances were small for all such comparisons. This confirms what can be seen via inspection of single-neuron examples: SMA responses were often similar, across cycling distances, when viewed aligned to movement end (Fig 2d-f).

In summary, SMA activity differs across steady-state cycles, even though muscle activity and M1 activity remain similar. This response specificity in SMA resembles, in some ways, contingency-specific activity in during movement sequences^19^ (e.g., a neuron that bursts only when pulling will be followed by turning). Yet specificity during cycling is manifested rather differently, by responses that evolve continuously, rather than burst at a key moment. The ramping activity we observed was more reminiscent of pre-movement responses in a timing task^20^. That said, ramping activity was not the only source of cycle-to-cycle response differences. To further explore the continuous unfolding of activity during cycling, we consider the evolution of population trajectories.

### SMA and M1 display different population trajectories

To gain intuition, we first visualize population trajectories in three dimensions (subsequent analyses will consider more dimensions). Projections onto the top three PCs are shown for one seven-cycle condition for M1 (Fig. 4a,b) and SMA (Fig. 4c,d). Traces are shaded light to dark to denote the passage of time. For the M1 populations, trajectories exited a baseline state just before movement onset, entered a periodic orbit during steady-state cycling, and remained there until settling back to baseline as movement ended. To examine within-cycle structure, we also applied PCA separately for each cycle (bottom of each panel). For M1, this revealed little new; the dominant structure on each cycle was an ellipse, in agreement with what was seen in the projection of the full response.

**Figure 4.**
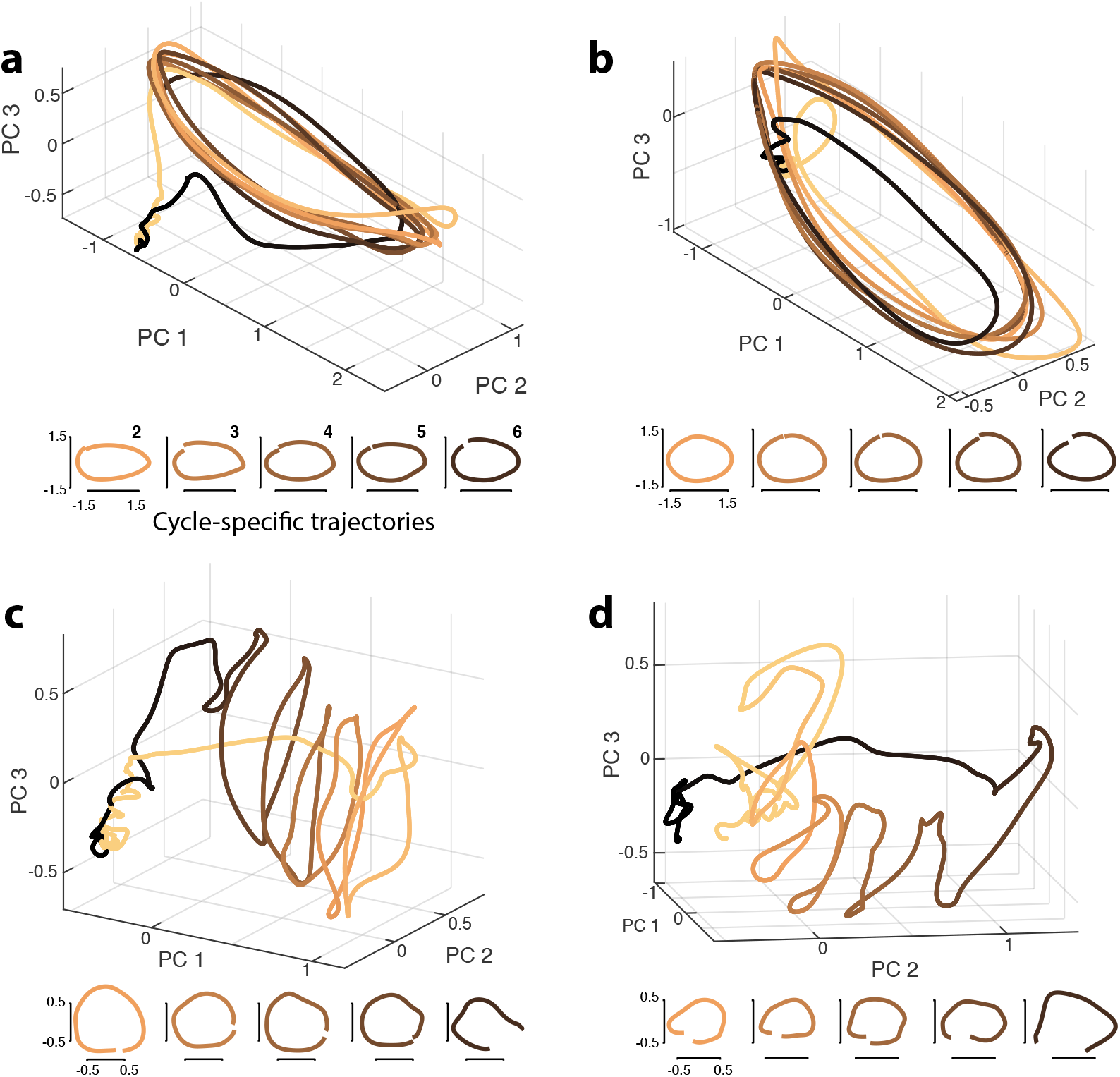
Visualization of population trajectories. **a)** M1 population trajectory corresponding during a seven-cycle movement (cycling forward from the bottom). The trajectory at top is the projection onto the top three PCs, and is shown from 1500 ms before movement onset until 500 after. The trajectory is shaded from tan (movement beginning) to black (movement end). PCs were found using all four seven-cycle conditions, using data from 200 ms before movement onset until 200 ms after movement ended (narrower than the plotted range, to prioritize dimensions that capture movement-related activity). Small plots at bottom show projections of each steady-state cycle (~500 ms of data) onto the first two PCs, founding using data from that cycle alone (with different PCs used for each steady-state cycle). This reveals the dominant structure on that cycle. Labels indicate cycle number (2-6). **b)** Same for monkey D, data corresponds to seven-cycle forward, top-start condition. **c)** SMA population trajectory for monkey C, corresponding to the same condition as in panel a. Analysis and plotting conventions are the same as for the analysis of M1. **d)** SMA population trajectory for monkey D, corresponding to the same condition as in panel b. Analysis and plotting conventions are the same as for the analysis of M1.

In SMA, the dominant geometry was quite different and also more difficult to summarize in three dimensions. We first consider data for monkey C (Fig. 4c). Just before movement onset, the population trajectory moved sharply away from baseline (from left to right in the plot). The trajectory then returned to baseline in a rough spiral, with each cycle separated from the last. The population trajectory for monkey D was different in some details (Fig. 4d) but it was again the case that a translation separated cycle-specific features.

SMA population trajectories appear to have a ‘messier’ geometry than M1 trajectories. In particular, cycle-specific loops appear non-elliptical and kinked. Yet it should be stressed that a three-dimensional projection is necessarily a compromise. The view is optimized to capture the largest features in the data; smaller features can be missed or partially captured and distorted. We thus employed cycle-specific PCs to visualize the shape of the trajectory on each cycle separately. Doing so revealed near-circular trajectories, much as in M1. Thus, individual-cycle orbits are present in SMA, but are a smaller feature relative to the large translation.

In summary, M1 trajectories are dominated by a repeating elliptical orbit while SMA trajectories are better described as helical. Each cycle involves an orbit, but these are separated by a translation. Also, unlike an idealized helix, individual-cycle orbits in SMA occur in somewhat different subspaces. This property is further explored and documented below.

### The SMA population response occupies different dimensions across cycles

We applied PCA separately for each cycle and computed ‘subspace overlap’: how well PCs derived from one cycle capture trajectories for the other cycles. For example, we computed PCs from the response during cycle one, projected the response during cycle two onto those PCs, and computed the percent variance explained. This was repeated across all cycle combinations. We employed six PCs, which captured most of the response variance for a given cycle. Essentially identical results were obtained using more PCs (Supp. Fig. 1). Variance was normalized so that unity indicates that two cycles occupy the same subspace. For comparison, we also analyzed muscle and M1 trajectories. As in Figure 3c-f, we compared within seven-cycle movements (Fig. 5a,c) and between seven- and four-cycle movements (Fig. 5b,d).

**Figure 5.**
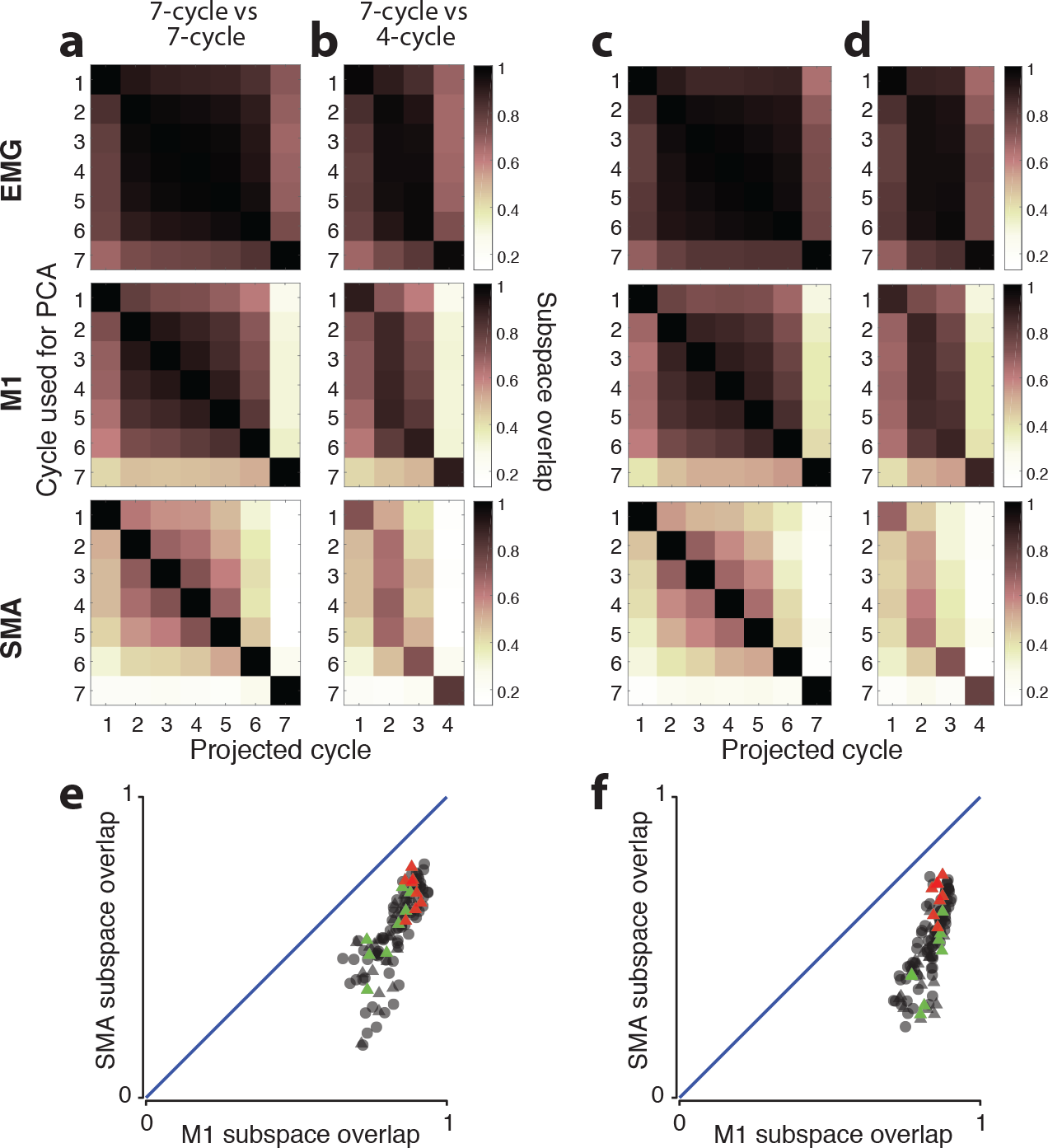
Subspace overlap between responses on different cycles. **a)** Matrices of subspace overlap when comparing cycles with a seven-cycle movement. Each matrix entry shows subspace overlap for one comparison. Matrix rows indicate the cycle used to find the PCs, and matrix columns indicate the cycle for which the variance captured is computed. For example, the entry in the second column / first row was found by projecting the population response for the second cycle onto the PCs found based on the first cycle. Data are averaged across conditions (two pedaling directions and two starting positions). Data are for monkey C. Diagonal entries necessarily have unity values, but matrices are not enforced to be symmetric (the subspace overlap measure is not symmetric). **b)** Subspace overlap when comparing between seven-cycle and four-cycle movements. Data are for monkey C. **c)** Same as panel a but for monkey D. **d)** Same as panel b but for monkey D. **e)** Subspace overlap for SMA versus M1 for all comparisons among steady-state cycles, both within seven-cycle movements (circles) and between seven-cycle and four-cycle movements (triangles). Each symbol corresponds to one comparison; e.g., the third cycle of a four-cycle movement with the fourth cycle of a seven-cycle movement. Data for each condition (different cycling directions and starting positions) are plotted separately. For each of the four conditions there are twenty within-seven-cycle comparisons (every steady-state cycle with every other steady-state cycle, excluding itself) and ten seven-versus-four-cycle comparisons. Red triangles highlight comparisons between cycles equidistant from movement end: six-of-seven versus three-of-four and five-of-seven versus two-of-four. Green triangles highlight comparisons between cycles equidistant from movement beginning: two-of-seven versus two-of-four and three-of-seven versus three-of-four. Data are for monkey C. **f)** Same for monkey D.

For the muscles, subspace overlap was high for all comparisons (Fig. 5a-d, top row). Subspace overlap was lower for M1 (middle row) yet still high. In particular, overlap was high among steady-state cycles, resulting in a central block structure. The block structure reveals that the subspace found for any of the steady-state cycles overlaps heavily with that for all the other steady-state cycles. For SMA, the central block was largely absent (bottom row). Comparing SMA versus M1, the average subspace overlap among steady-state cycles was 0.56 versus 0.83 (monkey C, p<0.0001 via bootstrap) and 0.51 versus 0.84 (monkey D, p<0.0001). Note that the changing subspace in SMA is not a consequence of the translating trajectory (Fig. 4); translation alters where activity is centered, not the subspace in which it resides.

For all comparisons among steady-state-cycles, across all conditions, subspace overlap was always lower for SMA versus M1 (Fig. 5e,f, p<10^−10^ via paired t-test for each monkey). Yet as with response distance, the magnitude of this effect depended on the situation. For example, subspace overlap in SMA tended to be higher when comparing cycles equidistant from movement’s end (red triangles) – e.g., cycle six-of-seven versus three-of-four. This effect can also be observed in Figure 5b,d (bottom): overlap is higher for the three-element diagonal ending in the bottom-right corner, relative to the diagonal starting in the top-left corner (p < 0.05 and p<.001 for monkey C and D, via bootstrap). This asymmetry was significantly greater in SMA versus M1 (p<0.05 for each monkey, via bootstrap).

### Population trajectories adopted by artificial networks

Our guiding hypothesis is that the SMA population response is structured to consistently differentiate between situations that will have different future motor outputs, even if the present motor output is the same. Such structure would be consistent with the idea that SMA internally tracks ‘motor context’ for the purpose of guiding future action. Consistent with this hypothesis, SMA activity differs across cycles, and occupies different subspaces across cycles. Intriguingly, SMA activity shows the least selectivity when there is no need to differentiate between situations; e.g., between cycle six-of-seven and cycle three-of-four, which lead to nearly identical future actions.

Yet is the roughly helical structure of the SMA population trajectory a natural solution when a network must track motor context? Are the properties documented above sufficient to ensure that SMA activity could consistently track context across times and conditions? To address these questions, we determine the critical properties of population trajectories displayed by simplified network models, and test whether those properties are present in the SMA population response.

For practical purposes, we define motor context as information that is important for guiding future movement but may not impact present motor output. Contextual information may be remembered (e.g., “I am performing a particular sequence”)^19^, internally estimated (“it has been 800 ms since the last button press”)^21^, or derived from abstract cues (“this fixation-point color means I must reach quickly when the target appears”)^42^. In the cycling task, salient contextual information arrives when the target appears, specifying the number of cycles to be produced. The current motor context (how many cycles remain) can then be updated throughout the movement, based on both visual cues and internal knowledge of the number of cycles already produced. Monkeys successfully used this contextual information; they essentially never stopped a cycle early or late. To ask how contextual information might be reflected in population trajectories, we trained artificial recurrent networks that did, or did not, need to internally track motor context.

We considered simplified inputs (pulses at specific times) and simplified outputs (pure sinusoids lasting four or seven cycles). We trained two families of recurrent networks. A family of ‘context-naïve’ networks received one input pulse, indicating that output generation should begin, and a different input pulse, indicating that output should be terminated. Initiating and terminating inputs were separated by four or seven cycles, corresponding to the desired output. Thus, context-naïve networks had no information regarding context until the arrival of the second input. Similarly, context-naïve networks had no need to track context; the key information was provided at the critical moment. A family of ‘context-tracking’ networks received only an initiating input. For context-tracking networks only, this input pulse differed depending on whether a four- or seven-cycle output should be produced. Context-tracking networks then had to generate a sinusoid with the appropriate number of cycles, and terminate appropriately with no further external guidance. For each family, we trained 500 networks that differed in their initial connection weights (*Methods*).

The two network families learned qualitatively different solutions involving population trajectories with different geometries. Context-naïve networks employed an elliptical limit cycle (Fig. 6a). The initiating input caused the network trajectory to enter an orbit, and the terminating input prompted the trajectory to return to baseline. This solution was not enforced but emerged naturally. There was network-to-network variation in how quickly activity settled into the limit cycle (Supp. Fig 2) but essentially all networks that succeeded in performing the task employed a version of this strategy.

**Figure 6.**
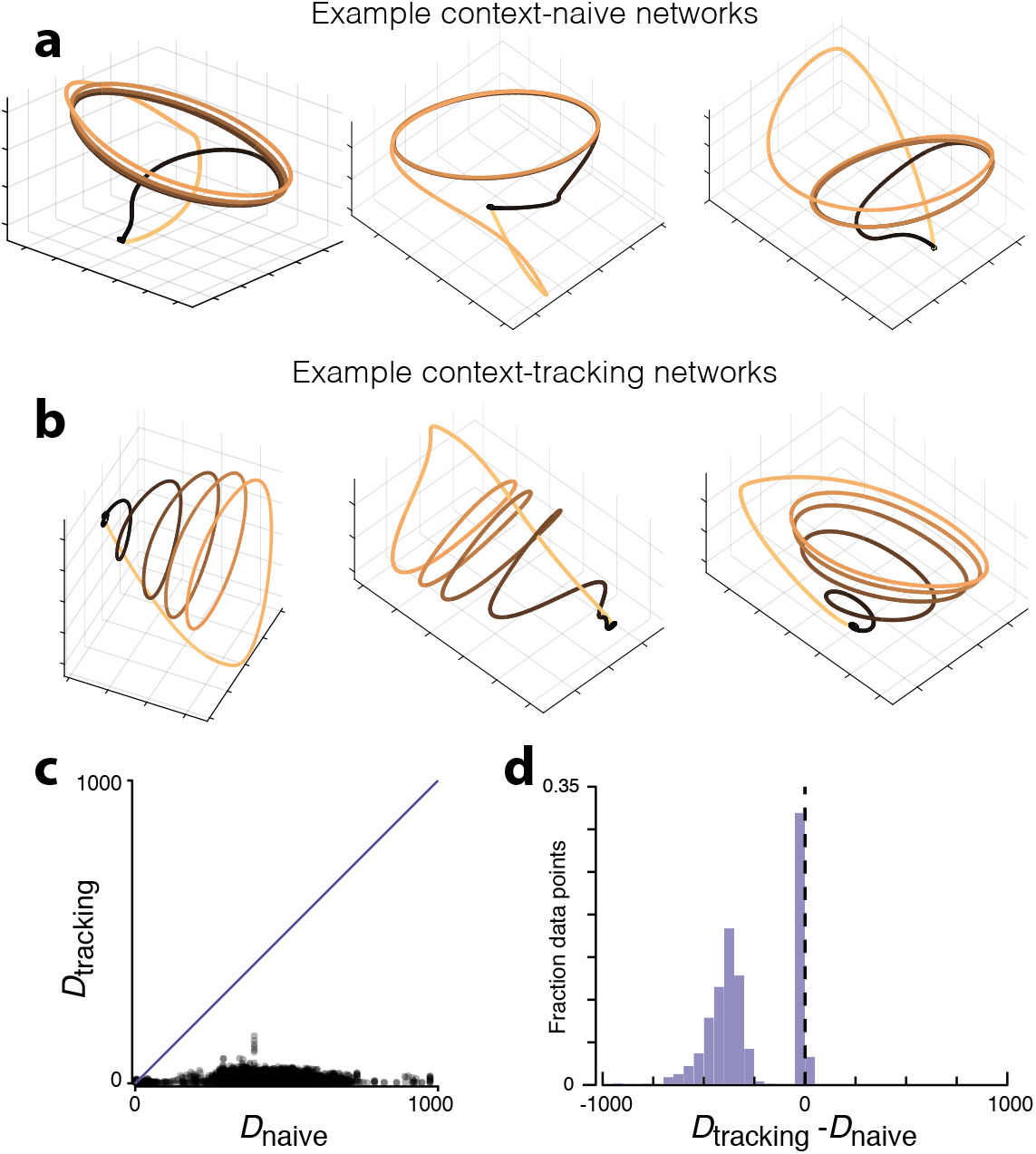
Trajectory geometry and divergence in simulated networks. **a)** Population trajectories for three example context-naïve networks during the four-cycle condition. For all examples, lower-left, lower-right, and vertical axes correspond to PC1, PC2, and PC3 respectively. **b)** Population trajectories for three example context-tracking networks. **c)** Trajectory divergence for context-tracking networks versus context-naïve networks. Comparison involves 500 networks of each type, paired arbitrarily. Each dot plots D_tracking_ versus D_naïve_ for one time during one pairing. Blue line indicates unity. **d)** Distribution of the differences in trajectory divergence between context-naïve and context tracking networks. Same data as in panel c, but for each time / network pair we computed D_tracking_ – D_naïve_.

Context-tracking networks utilized population trajectories that were more helical; the trajectory on each cycle was separated from the others by an overall translation (Fig. 6b). While there was network-to-network variability in the exact learned trajectory (Supp. Fig 3), all successful context-tracking networks employed some form of helical or spiral trajectory. This solution is intuitive: context-tracking networks do not have the luxury of following a repeating orbit. If they did, information regarding context would be lost, and the network would have no way of ‘knowing’ when to cease producing the output.

For context-tracking networks, trajectories could also occupy somewhat different subspaces on different cycles. Projected onto three dimensions, this geometry resulted in individual-cycle trajectories of seemingly different magnitude (first and third examples in Fig. 6b). As with the helical structure, this geometry creates separation between individual-cycle trajectories. There was considerable variation in the degree to which this strategy was employed. Some context-tracking networks used nearly identical subspaces for every cycle while others used quite different subspaces. Context-naïve networks never employed this strategy; the same limit cycle was always followed across steady-state cycles.

The population geometry adopted by context-naïve and context-tracking networks bears obvious similarities to the empirical population geometry in M1 and SMA, respectively. That said, we stress that neither family is intended to faithfully model the corresponding area. Furthermore, a number of reasonable alternative modeling choices exist. For example, rather than asking context-tracking networks to track progress using internal dynamics alone, one can provide a ramping input that does so. Interestingly, context-tracking networks trained in the presence / absence of ramps employed very similar population trajectories (Supp Fig 4). The slow translation that produces helical structure is a useful computational tool – one that networks produced on their own if needed but were also content to inherit from upstream sources. For these reasons, we focus not on the details of specific network trajectories, but rather on the geometric features that differentiate context-tracking from context-naïve network trajectories, and that might similarly differentiate M1 and SMA population trajectories.

### Trajectory divergence

The trajectories displayed by context-tracking networks reflect specific solutions to a general problem: ensuring that two trajectory segments never trace the same path and then diverge. Avoiding such divergence is critical when network activity must distinguish between situations that have the same present motor output, but different future outputs. Rather than assessing the specific path of particular solutions (which differed across networks), we developed a general metric of trajectory divergence. We note that trajectory divergence differs from trajectory tangling^38^, which was very low in both SMA and M1 (Supp Fig 5). Trajectory tangling assesses whether trajectories are consistent with a locally smooth flow-field. Trajectory divergence assesses whether similar paths eventually separate, smoothly or otherwise. A trajectory can have low tangling but high divergence, or vice versa (Supp Fig 6).

To construct a metric of trajectory divergence, we consider times *t* and *t*′, associated population states ***x***_*t*_ and ***x***_*t*′_, and future population states ***x***_*t*+Δ_ and ***x***_*t*′+Δ_. We consider all possible pairings of *t* and *t*′. For example, *t* and *t*′ might occur during different cycles of the same movement or during different cycling distances. We compute the ratio 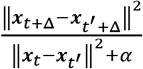, which becomes large if ***x***_*t*+Δ_ differs from ***x***_*t*′+Δ_ despite ***x***_*t*_ and ***x***_*t*′_ being similar. The constant *α* is small and proportional to the variance of ***x***, and prevents hyperbolic growth.

Given that the difference between two random states is typically sizeable, the above ratio will be small for most values of *t*′. As we are interested in whether the ratio ever becomes large, we take the maximum, and define divergence for time *t* as:

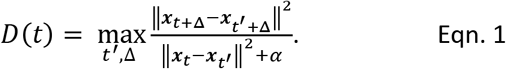

We consider only positive values of Δ. Thus, *D*(*t*) becomes large if similar trajectories diverge but not if dissimilar trajectories converge. Divergence was assessed using a twelve-dimensional neural state. Results were similar for all reasonable choices of dimensionality.

*D*(*t*) differentiated between context-tracking and context-naïve networks. We compared these two classes by considering pairs of networks, one context-tracking and one context-naïve. For each time, we plotted *D*(*t*) for the context-tracking network versus that for the context-naïve network. Trajectory divergence was consistently lower for context-tracking networks (Fig. 6c, p<0.0001, rank sum test). This was further confirmed by considering the difference in the values of *D*(*t*), for all times and all network pairs (Fig. 6d). Both context-tracking and context-naïve trajectories contained many moments when divergence was low, resulting in a narrow peak near zero. However, context-naïve trajectories (but not context-tracking trajectories) also contained moments when divergence was high, yielding a large set of negative differences.

### Trajectory divergence is lowest for SMA

The roughly helical structure of the empirical SMA population response (Fig. 4) suggests low trajectory divergence, as does the finding that SMA responses differ across cycles (Figs. 3 and 5). Yet the complex shape of the empirical trajectories makes it impossible to ascertain, via inspection, whether divergence is low. Furthermore, it is unclear whether cycle-to-cycle response differences consistently ensure low divergence across times and all cycling distances. We therefore directly measured trajectory divergence for the empirical trajectories.

Plotting SMA versus M1 trajectory divergence for each time (Fig. 7a,b) revealed that divergence was almost always lower in SMA. We next computed distributions of the difference in divergence, at matched times, between SMA and M1 (Fig. 7c,d). There was a narrow peak at zero (times where divergence was low for both) and a large set of negative values, indicating lower divergence for SMA. Strongly positive values (lower divergence for M1) were absent (monkey C) or very rare (monkey D; 0.13% of points > 20). Via bootstrap, distributions were significantly negative for both monkeys (p<0.00001 for each). It was also the case that trajectory divergence was much lower in SMA than in the muscle populations (Supp Fig 7). The overall scale of divergence values was smaller for the empirical data versus the networks. Specifically, divergence reached higher values for context-naïve networks than for the empirical M1 trajectories. This occurs because simulated trajectories can repeat almost perfectly, yielding very small values of the denominator in equation 1. Other than this difference in scale, trajectory divergence for SMA and M1 differed in much the same way as for context-tracking and context-naïve networks (compare Fig. 7c,d with Fig. 6d).

**Figure 7.**
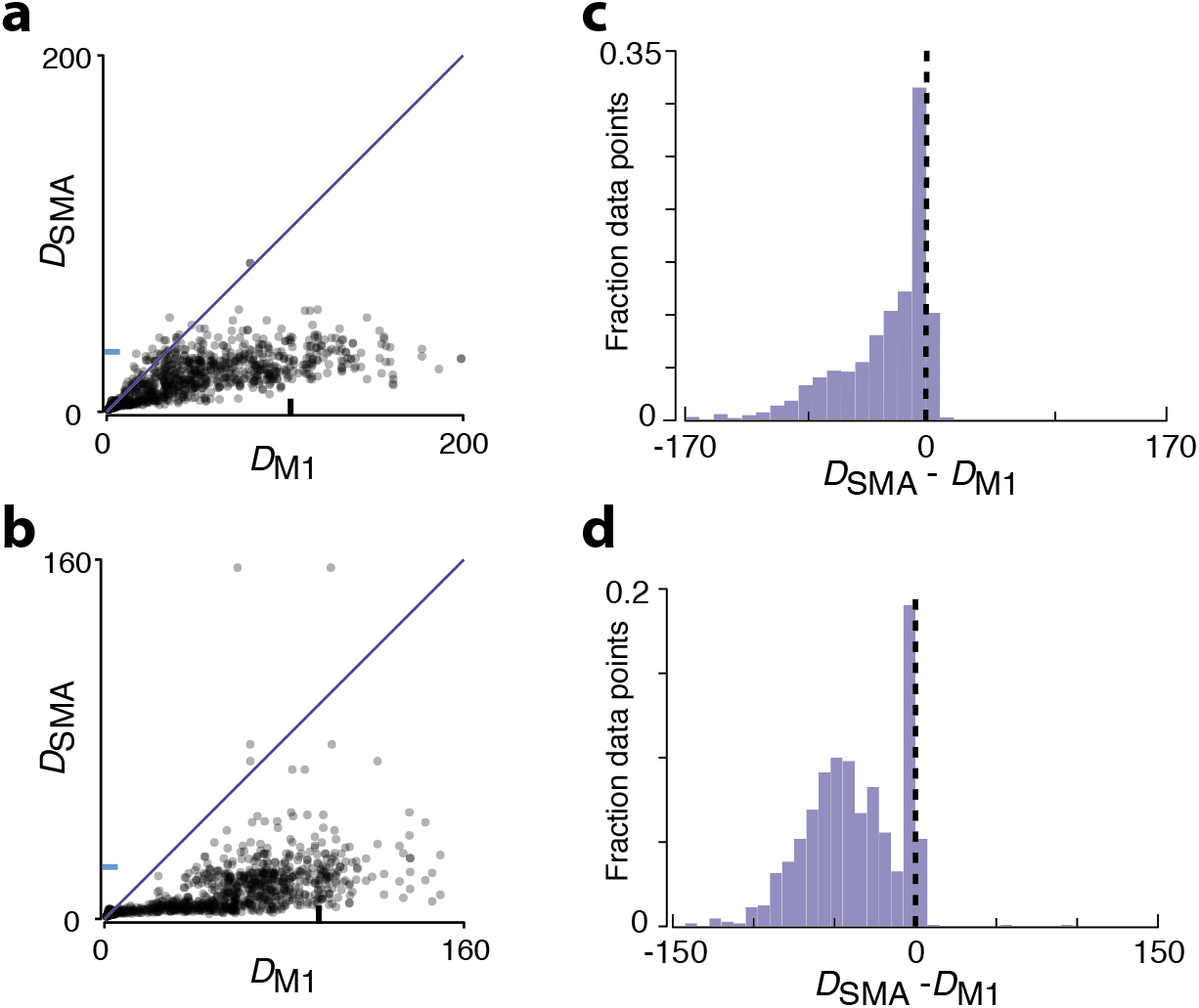
Trajectory divergence in M1 and SMA. **a)** Trajectory divergence for SMA versus M1 (monkey C). Each dot corresponds to one time during one condition. Divergence was computed considering all times for all conditions within a given starting position / pedaling direction. Data for all conditions is then plotted together. Blue tick mark on the vertical axis denotes 90th percentile trajectory divergence for SMA. Black tick mark along the horizontal axis denotes 90th percentile trajectory divergence for M1. **b)** Same for monkey D **c)** Distribution of the differences in trajectory divergence between SMA and M1 for monkey C. Same data as in panel a, but for each time / condition we computed the difference in divergence. **d)** Same for monkey D

The ability to consider both network and neural trajectories (despite differences across networks and across monkeys) underscores that the divergence metric describes trajectory geometry at a useful level of abstraction. Multiple specific features can contribute to low divergence, including ramping activity, cycle-specific responses, and the use of different subspaces on different cycles. Different network instantiations may use these different ‘strategies’ to different degrees. Trajectory divergence provides a useful summary of a computationally relevant property, regardless of the specifics of how it was achieved.

Because trajectory divergence abstracts away from the details of specific trajectories, it can be readily applied in new situations. For example, the cycling task involved not only different cycling distances, but also different cycling directions and different starting positions. The latter is particularly relevant, because movements ended at the same position (top versus bottom of the cycle) as they started. Thus, how a movement will end depends on information present at the movement’s beginning. One could ask whether SMA responses keep track of such information by assessing ‘starting-position-tuning’ in a variety of ways, following the example of Figures 3 and 5. However, it is simpler, and more relevant to the hypothesis being considered, to ask whether divergence remains low when comparisons are made across all conditions including starting positions. This was indeed the case (Supp Fig 8).

### Computational implications of trajectory divergence

We considered trajectory divergence because of its expected computational implications. A network with a high-divergence trajectory can accurately and robustly generate its output on short timescales. Yet unless guided by external inputs at key moments, such a network may be susceptible to errors on longer timescales. For example, if a trajectory approximately repeats, a likely error would be the generation of extra cycles or the inappropriate skipping of a cycle.

To test whether these intuitions are accurate, we performed additional simulations. We employed an atypical training approach that enforced an internal network trajectory, as opposed to the usual approach of training a target output. We trained networks to precisely follow the M1 trajectory, recorded during a four-cycle movement, without any input indicating when to stop (Fig. 8a). To ensure that the solutions found were not overly delicate, networks were trained in the presence of additive noise. Using data from each monkey, we trained forty networks: ten for each of the four four-cycle conditions. Networks were able to reproduce the cyclic portion of the M1 trajectory. However, without the benefit of a stopping pulse, networks failed to consistently follow the end of the trajectory. For example, networks sometimes erroneously produced extra cycles (Fig. 8b) or skipped cycles and stopped early (Fig. 8c).

**Figure 8.**
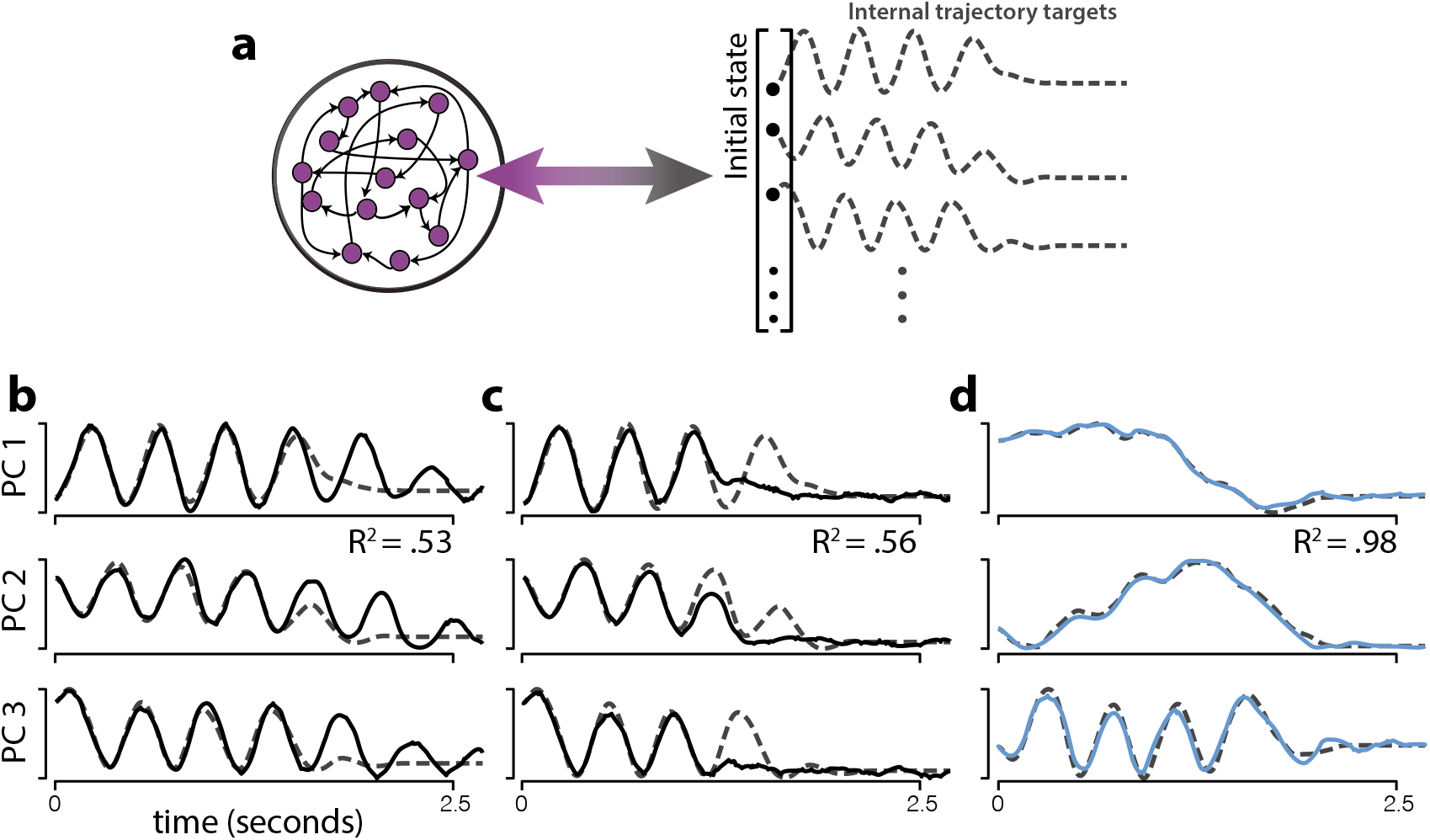
Example behavior of networks trained to follow the empirical M1 or SMA population trajectory. **a)** Illustration of the trajectory-constrained neural networks. Networks were trained to autonomously follow a target trajectory defined by the top six PCs of the empirical population trajectory, during a four-cycle movement, from movement onset until 250ms after movement offset. Dashed lines show the target trajectory for three PCs for one example: the M1 trajectory for monkey D, cycling backward starting at the bottom. The activity of every neuron in the network was trained to follow a random combination of the projection onto the top six PCs. This ensured that the simulated population trajectory matched the empirical trajectory. **b)** Example target (dashed) and produced (solid) network trajectories on one trial, after training is complete. Target trajectory was the empirical M1 trajectory. The trajectory produced by the network initially matches the target, but continues ‘cycling’ past when it should have ended. This resulted in an R2 (variance in the target accounted for by the produced trajectory) considerably below unity. **c)** As in panel b, but for an example trial where the opposite error was made: the network trajectory stops cycling earlier than it should have. This trajectory is produced by the same network as in b, and is attempting to match the same target. The only difference is the additive noise on that particular trial. **d)** Example target (dashed) and produced (blue) network trajectories on one trial for a network trained to produce the empirical SMA trajectory. The level of additive noise was the same as for the network in panels b and c, but the network does not fail to follow the trajectory to the end.

We also trained networks to follow the empirical SMA trajectories. Those trajectories contained both a rhythmic component and lower-frequency ‘ramping’ signals (Figure 8d) related to the translation visible in Fig. 4c,d. In contrast to the high-divergence M1 trajectories, which were never consistently followed for the full trajectory, the majority of network initializations resulted in good solutions where the low-divergence SMA trajectory was successfully followed from beginning to end. Thus, in the absence of a stopping pulse, the empirical SMA trajectories could be produced and could terminate reliably in a way that the empirical M1 trajectories could not.

## Discussion

Prior studies argue that SMA contributes to the guidance of action based on internal, abstract or contextual factors^7,11,14–17,19,20^. We translated this conceptual hypothesis into a prediction regarding the geometry of population activity: trajectory divergence should be consistently low. This hypothesis embodies an essential component of prior ideas. The ability to internally guide action implies that activity should be structured to reliably differentiate between situations that are the same now but will soon become different. We tested whether low trajectory divergence was observed in a novel task, whether low divergence is shared between network models and empirical data, and whether low divergence might provide a cohesive explanation for diverse features of neural responses.

We employed our recently developed cycling task both because it has proved useful in characterizing population geometry in M1^38^, and because it produces multiple instances of behavioral divergence: situations with the same current motor output but different future motor outputs. The cycling task is neither a sequence task nor a timing task, yet it shares commonalities with both paradigms. Consistent with this, there were both differences and commonalities in single-neuron response features during cycling and during other tasks. The ramping firing rates we observed resemble those seen in timing tasks^20^. We also observed cycle-specific responses – e.g., different firing rates across repeated cycles – which may be thought of as a form of sequence selectivity. However, cycle-selectivity was produced not by response bursts tied to a particular contingency^19^, but by a combination of ramping and cyclic activity, with different subspaces being occupied on different cycles. Selectivity for cycling distance (e.g., different responses when starting a four-versus seven-cycle movement) can also be seen as related to sequence selectivity. Yet such selectivity was not equally present across all comparisons; it was pronounced when comparing situations where future motor output would be different.

These diverse response features can be understood in a unified way: they all serve to reduce divergence. The resulting shape of the SMA population trajectory was a rough helix. This can be seen as a neural ‘strategy’ for avoiding trajectory divergence by reliably differentiating among situations that have the same present motor output but different future outputs. In contrast, M1 population trajectories traced out elliptical orbits and had high trajectory divergence.

Simulations confirmed that divergence was naturally high in networks that did not have to internally track context; context-naïve networks displayed elliptical population trajectories resembling the dominant structure in M1. Conversely, divergence was low in networks that had to track context. Context-tracking networks displayed helical population trajectories that resembled simplified SMA trajectories. Although some kind of helical structure was universal across such networks, there was variability in the exact solution. For some networks, low-divergence was achieved solely through the translation that separated cycles, while in other networks different cycles occupied somewhat different subspaces (as in the neural data, but typically to a lesser degree). This underscores the value of a metric such as trajectory divergence, which can abstract away from solution-specific features and summarize whether a trajectory is appropriate for a type of computation.

The present study builds upon recent studies examining the shape and nature of population activity to evaluate hypotheses regarding the network-level computation. Most such studies quantify specific features that relate to how a network might perform the task of interest^15,30,37,45–48^, and this will remain an essential strategy. Yet, as noted above, one may also wish metrics of population geometry that are more general, and quantify properties that may be preserved across a class of computations regardless of the particular instantiation. Our divergence metric was designed with this goal in mind. As another example, we recently characterized a different geometric property, trajectory tangling, when examining the M1 population response^38^. Low tangling is necessary for a network to robustly generate an output via internal dynamics. We found that trajectory tangling was much lower for M1 trajectories than for muscle population trajectories^38^. That difference was apparent across tasks and species, and helped explain seemingly paradoxical features of M1 activity. The presence of low tangling in M1, but not sensory areas, argued that M1 activity is shaped by the need to perform robust pattern generation. In the present study, we found that trajectory tangling was similarly low in both SMA and M1, consistent with activity in both areas being strongly shaped by internal dynamics that need to provide a temporally structured output. However, the nature of the computation performed by those internal dynamics is likely very different. Only in SMA do population trajectories show low trajectory divergence, consistent with guidance of movement based on contextual information.

The property of low-divergence might help explain the diversity of SMA response properties both within and between tasks. In particular, low divergence would be consistent with single-neuron response properties such as sequence-selectivity (though whether such selectivity actually leads to consistently low divergence still needs to be confirmed). Conversely, there may be situations where divergence becomes high in SMA in a revealing way. For example, there are likely limits on the timescales across which SMA (or the monkey as a whole) can internally track context, and this might be revealed in the timescales over which divergence stays low. For example, it seems unlikely that cycling for one hundred cycles would be accompanied by an extended low-divergence helix with one hundred separated loops.

As another example, trajectory divergence is unlikely to remain low when action is guided by sudden, unpredictable cues. These predictions are testable, and underscore that the expectation of low trajectory divergence is not universal. Trajectory divergence is expected to be low only when SMA is in fact consistently tracking context. This yields predictions that depend, in a useful way, on hypothesized role of SMA for the task being studied.

A reasonable question is why low trajectory divergence is not a more universal property of movement-related population activity. Why not employ a fully unified strategy, where a low-divergence trajectory produces the final motor output (as occurs in our simple context-tracking networks)? Why have separate areas – SMA and M1 – with low and high trajectory divergence? The presence of a high-divergence area may be useful for two reasons. One is that dispensing with divergence-avoiding signals yields greater dynamic range available for generating the fine features of the motor output. A second is that a repeating trajectory may allow adaptation that occurs on one cycle to naturally generalize to other cycles. Thus, the degree of trajectory divergence in a given area has implications regarding what types of learning are possible, and may indicate the ‘level’ of control provided by a given motor area.

## STAR Methods

### CONTACT FOR REAGENT AND RESOURCE SHARING

Further information and requests for resources and reagents should be directed to and will be fulfilled by the Lead Contact, Dr. Mark M. Churchland.

### EXPERIMENTAL MODEL AND SUBJECT DETAILS

#### Main experimental datasets

Subjects were two adult male rhesus macaques (monkeys C and D). Animal protocols were approved by the Columbia University Institutional Animal Care and Use Committee. Experiments were controlled and data collected under computer control (Speedgoat Real-time Target Machine). During experiments, monkeys sat in a customized chair with the head restrained via a surgical implant. Stimuli were displayed on a monitor in front of the monkey. A tube dispensed juice rewards. The left arm was loosely restrained using a tube and a cloth sling. With their right arm, monkeys manipulated a pedal-like device. The device consisted of a cylindrical rotating grip (the pedal), attached to a crank-arm, which rotated upon a main axel. That axel was connected to a motor and a rotary encoder that reported angular position with 1/8000 cycle precision. In real time, information about angular position and its derivatives was used to provide virtual mass and viscosity, with the desired forces delivered by the motor. The delay between encoder measurement and force production was 1 ms.

Horizontal and vertical hand position were computed based on angular position and the length of the crank-arm (64 mm). To minimize extraneous movement, the right wrist rested in a brace attached to the hand pedal. The motion of the pedal was thus almost entirely driven by the shoulder and elbow, with the wrist moving only slightly to maintain a comfortable posture.

### METHOD DETAILS

#### Task

Monkeys performed the ‘cycling task’ as described previously^38^. The monitor displayed a virtual landscape, generated by the Unity engine (Unity Technologies, San Francisco). Surface texture and landmarks provided visual cues regarding movement through the landscape along a linear ‘track’. One rotation of the pedal produced one arbitrary unit of movement. Targets on the track indicated where the monkey should stop for juice reward.

Each trial of the task began with the appearance of an initial target. Each trial began with the monkey stationary on top of an initial target. After a 1000 ms hold period, the final target appeared at a prescribed distance. Following a randomized (500-1000 ms) delay period, a go-cue (brightening of the final target) was given. The monkey then had to cycle to acquire the final target. After remaining stationary in the final target for 1500 ms, the monkey received a reward.

The full task included 20 conditions distinguishable by final target distance (one-, two-, four-, and seven-cycles), initial starting position (top or bottom of the cycle), and cycling direction (forward or backward). Half-cycle distances were also included in the task (evoking quite brief movements). Because of the absence of a full-cycle response, they are not amenable to many of the analyses we employ, and were thus not analyzed.

Salient visual cues (landscape color) indicated whether cycling must be ‘forward’ (the hand moved away from the body at the top of the cycle) or ‘backward’ (the hand moved toward the body at the top of the cycle) to produce forward virtual progress. Trials were blocked into forward and backward cycling. Other trials types were interleaved using a block-randomized design. For each neural / muscle recording, we collected a median of 15 trials / condition for both monkeys.

#### Neural recordings during cycling

After initial training, we performed a sterile surgery during which monkeys were implanted with a head restraint and recording cylinders (Crist Instruments, Hagerstown, MD). For M1 recordings, cylinders were placed surface normal to the cortex and centered over the border between caudal PMd and primary motor cortex. After recording in M1, we performed a second sterile surgery to move the cylinders over the SMA. SMA cylinders were located over the SMA as determined from a previous magnetic resonance imaging scan, and were angled at ~20° degrees to avoid the central sulcus vein. The skull within the cylinders was left intact and covered with a thin layer of dental acrylic. Electrodes were introduced through small (3.5 mm diameter) burr holes drilled by hand through the acrylic and skull, under ketamine / xylazine anesthesia. Neural recordings were made using conventional single electrodes (Frederick Haer Company, Bowdoinham, ME) driven by a hydraulic microdrive (David Kopf Instruments, Tujunga, CA). The use of conventional electrodes, as opposed to electrode arrays, allowed recordings to be made from the medial bank (where most of the SMA is located) and from both surface and sulcal M1.

Recording locations were guided via microstimulation, light touch, and muscle palpation protocols to confirm the trademark properties of each region. For motor cortex, recordings were made from primary motor cortex (both surface and sulcal) and the adjacent (caudal) aspect of dorsal premotor cortex. These recordings are analyzed together as a single motor cortex population. All recordings were restricted to regions where microstimulation elicited responses in shoulder and arm muscles.

Neural signals were amplified, filtered, and manually sorted using Blackrock Microsystems hardware (Digital Hub and 128-channel Neural Signal Processor). A total of 380 isolations were made across the two monkeys. On each trial, the spikes of the recorded neuron were filtered with a Gaussian (25 ms standard deviation; SD) to produce an estimate of firing rate versus time. These were then temporally aligned and averaged across trials as previously described^38^. Averages were made across a median of 15 trials/condition.

#### EMG recordings

Intra-muscular EMG was recorded from the major shoulder and arm muscles using percutaneous pairs of hook-wire electrodes (30mm x 27 gauge, Natus Neurology) inserted ~1 cm into the belly of the muscle for the duration of single recording sessions. Electrode voltages were amplified, bandpass filtered (10-500 Hz) and digitized at 1000 Hz. To ensure that recordings were of high quality, signals were visualized on an oscilloscope throughout the duration of the recording session. Recordings were aborted if they contained significant movement artifact or weak signal. That muscle was then re-recorded later. Offline, EMG records were high-pass filtered at 40 Hz and rectified. Finally, EMG records were smoothed with a Gaussian (25 ms SD, same as neural data) and trial averaged (see below). Recordings were made from the following muscles: the three heads of the *deltoid*, the two heads of the *biceps brachii*, the three heads of the *triceps brachii, trapezius, latissimus dorsi, pectoralis, brachioradialis, extensor carpi ulnaris, extensor carpi radialis, flexor carpi ulnaris, flexor carpi radialis*, and *pronator*. Recordings were made from 1-8 muscles at a time, on separate days from neural recordings. We often made multiple recordings for a given muscle, especially those that we previously noted could display responses that vary with recording location (*e.g*., the *deltoid*).

### QUANTIFICATION AND STATISTICAL ANALYSIS

#### Preprocessing and PCA

Many of our analyses employ PCA, either as a denoising step or as an essential aspect of the analysis. Because PCA seeks to capture variance, it can be disproportionately influenced by differences in firing rate range (*e.g*., a neuron with a range of 100 spikes/s has 25 times the variance of a similar neuron with a range of 20 spikes/s). This concern is larger still for EMG, where the scale is arbitrary and can differ greatly between recordings. The response of each neuron / muscle was thus normalized prior to application of PCA. EMG data were fully normalized: *response*: = *response/range*(*response*), where the range is taken across all recorded times and conditions. Neural data were ‘soft’ normalized: *response*: = *response*/(*range*(*response*) + 5). We standardly^38,46,49^ use soft normalization to balance the desire for PCA to explain the responses of all neurons with the desire that weak responses not contribute on an equal footing with robust responses. Soft normalization is also helpful for other analyses (e.g,. of response distance) to avoid results being dominated by a few high-firing-rate neurons.

To perform PCA, neural data were formatted as a ‘full-dimensional’ matrix, *X^full^*, of size *n×t*, where *n* is the number of neurons (or muscles) and *t* indexes across the analyzed times and conditions. PCA was used to find the PCs, *V*, and a reduced-dimensional version of the data, *X*, such that *X^full^* ≈ *VX*, where *V* are the PCs (‘dimensions’ upon which the data are projected). The set of times and conditions considered varied by analyses. We always employed enough PCs to capture the majority of the data variance for all populations. For example, for analyses of divergence, we employed twelve PCs, which captured an average of 89% and 87% of the data variance in M1 and SMA respectively.

#### Response distance

Response distance assesses the degree to which the population response is different for the response on two different cycles (either within a seven-cycle movement, or between seven-cycle and four-cycle movements of the same type). Response distance was computed after using PCA to denoise responses. For each population, we replaced the recorded estimate of firing rate with a denoised estimate of firing rate reconstructed from the top twelve PCs. This use of PCA denoises the response of each neuron based on the commonalities present across the entire population.

Response distance – the root-mean-squared difference in the firing rate of each neuron – after denoising is equivalent to the distance in PCA space. Thus, for practical purposes response distance was computed in this manner. Results were virtual identical if PCA was not used to denoise the data, except that sampling error slightly inflated all response distances (due to sampling error, even cycles with truly identical responses would not show a zero response distance).

We also wished to ensure that response distance was not inflated if two cycles had similar responses but different durations. This is of little concern when comparing among steady-state cycles (duration was highly stereotyped) but becomes a concern when comparing an initial-cycle response with a steady-state cycle response. To avoid misalignment, the response on each cycle was scaled to have the same duration, with the angular position matched at all times. After alignment, response distance is zero if two responses are the same except for their time-course.

Comparisons were made within a given seven-cycle condition and between seven-cycle and four-cycle conditions. Comparisons were always made between conditions of the same type (i.e., the same cycling direction and starting position). Response distances were normalized by response magnitude (the root-mean-squared difference in firing rate from its mean) within a steady-state cycle of the same condition type. For simplicity, we chose the fourth cycle of the seven-cycle movement.

#### Subspace overlap

Subspace overlap was used to measure the degree to which the population response occupied different neural dimensions on different cycles (different cycles within a distance, or between distances).

Subspace overlap was always computed for a pair of cycles: a reference cycle and a comparison cycle. PCA was applied to the population response for the reference cycle, to obtain six cycle-specific PCs. These are the dimensions that best capture the response on that cycle, during that particular distance, cycling direction, and starting position. The population response for the comparison cycle was then projected onto those PCs, and the variance captured was computed. This variance was normalized by the variance captured if comparison-cycle data were projected onto ‘native’ PCs, computed from the comparison cycle population response. The resulting ‘subspace overlap’ is thus unity if the population response on the reference and comparison cycles occupies the same dimensions (i.e., are spanned by the same PCs).

To test for statistical significance, we used a bootstrap procedure. For each population, we resampled all neurons with replacement and repeated the analysis. Resampling was performed 1000 times. For analyses that compared SMA and M1, comparison was performed across all pairs of SMA and M1 bootstrapped datasets (1 million comparisons).

#### Trajectory Divergence

Consider times *t* and *t*′. These times could occur within the same movement. E.g., *t* could be a time near the middle of the movement and *t*′ could be a time near the end. The two times could also occur for different distances within the same condition type. E.g., if we consider forward cycling that starts at the top, *t* could occur during a two-cycle movement and *t*′ could occur during a seven-cycle movement. Consider the associated neural states ***x***_*t*_ and ***x***_*t*′_. The squared distance between these states is ||***x***_*t*_ − ***x***_*t*′_||^2^. The squared distance between the corresponding neural states, some time Δ in the future, is ||***x***_*t*+Δ_ − ***x***_*t*′+Δ_||^2^. Divergence assess whether this future distance ever becomes large despite the present distance being small. We define the divergence for a given time, during a given condition, as:

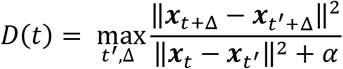

Where *t*′ indexes across all times within all movements of the same type, and Δ indexes from one to the largest time that can be considered: min(*T* − *t, T*′ − *t*′) where *T* is the duration of the condition associated with time *t* and *T*′ is the duration of the condition associated with time *t*′. The states ***x*** are rows from the matrix *X*, after application of PCA. PCA was applied to a matrix *X^full^* that contained data from 100 ms before movement onset until 100 ms after movement ended, for all movements of the type being considered (e.g., all four distances for forward cycling starting at the top). We employed a twelve-dimensional ***x*** (i.e., the projection onto the top twelve PCs). Results were not sensitive to the choice of dimensionality; divergence was always much lower for SMA versus M1. This was also true if we did not employ PCA at all, but simply used *X^full^*. That said, we still preferred to use PCA as a preprocessing step. Reducing dimensionality makes analysis much faster, and denoising reduces concerns about the denominator fluctuating due to sampling error. To ensure the denominator was well behaved (e.g., did not become too close to zero) we also included the constant *α*, set to 0.01 times the variance of *X*. Results were essentially identical across a range of reasonable values of *α*. For our primary analysis, divergence was measured separately for each of the four condition types. Thus, *t*′ indexes across times and across distances within the same condition type. The same effect held (SMA divergence lower than M1 divergence) if *t*′ indexed across all other conditions (Supp Fig 8).

#### Recurrent Neural Networks

We trained recurrent neural networks to produce four and seven cycles of a sinusoid in response to external inputs. A network consisted of *N* = 50 firing-rate units with dynamics:

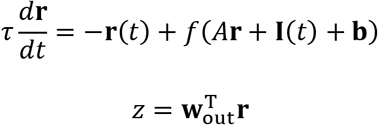

where *τ* is a time-constant, **r** represents an *N*-dimensional vector of firing rates, *f*: = tanh is a nonlinear input-output function, *A* is an *N* × *N* matrix of recurrent weights, **I**(*t*) represents time-varying external input, and **b** is a vector of constant biases. The network output *z* is a linear readout of the rates. Both *A* and **w**_out_ were initially drawn from a normal distribution of zero mean and variance 1/*N*. **b** was initialized to zero. Throughout training, *A*, **w**_out_, and **b** were modified.

Context-tracking networks were trained to generate a four-cycle versus seven-cycle output after receiving a short go pulse (a square pulse that lasts for half a cycle duration prior to the start of the output) without the benefit of a stopping pulse. For context-tracking networks only, go pulses were different depending on whether four or seven cycles should be produced. The two go pulses were temporally identical, but entered the network through different sets of random input weights; **I**(*t*) = **w**_4_*I*(*t*) or **I**(*t*) = **w**_7_*I*(*t*), where *I*(*t*) is a square pulse of unit amplitude.

Context-naïve networks received both a go pulse and a stop pulse. Go and stop pulses were distinguished by entering the network through different sets of random input weights; **I**(*t*) = **w**_go_*I*(*t*) or **I**(*t*) = **w**_stop_*I*(*t*). Go and stop pulses were separated by an appropriate amount of time to compete the desired number of cycles. We only analyzed outputs when go and stop pulses were separated by four or seven cycles. Yet we did not wish context-naïve networks to learn overly specific solutions. Thus, during training, we also included trials where the network had to cycle continuously in the absence of a stop-pulse. This ensured that context-naïve networks learned a general solution; e.g., could cycle for six cycles and stop if the go and stop pulses were separated by six cycles.

We also considered a modification of context-naïve networks, that received an external timing signal. Rather than a distinct ‘stop pulse’, modified-context-naïve networks received a downward ramping input through another set of weights **w**_ramp_. The ramping input has a constant slope but different starting values for different numbers of desired cycles. The end of the cycling period in this case was indicated by the ramp signal reaching zero.

In all three cases, networks were trained using back-propagation-through-time^50^ using TensorFlow and an Adam optimizer to adjust *A*, **w**_out_, and **b** to minimize the squared difference between the network output *z* and the sinusoidal target function. All the input weights, **w**_4_, **w**_7_, **w**_go_, **w**_stop_ and **w**_ramp_, were drawn from a zero-mean unit-variance normal distribution and remain fixed throughout training. The amplitude of pulses and cycles were set to a value that produced a response but avoided saturating the units. The height of the ramp signal was set to the same amplitude as the input pulses for the sevenendnot-cycle condition. For each condition, we trained 500 networks, each initialized with a different realization of *A* and **w**_out_.

#### Trajectory-constrained Neural Networks

To test the computational implications of trajectory divergence, we trained recurrent neural networks with an atypical approach. Rather than training networks to produce an output, we trained them to autonomously follow a target internal trajectory^38,51^. We then asked whether networks were able to follow those trajectories from beginning to end, without the benefit of any inputs indicating when to stop.

Target trajectories were derived from neural recordings (M1, and SMA) during the four-cycle movements for each of the four condition types (forward-bottom-start, forward-top-start, backward-bottom-start, backward-top-start). Target trajectories spanned the time period from movement onset until 250 ms after movement offset. To emphasize that the network should complete the trajectory and remain in the final state, we extended the final sample of the target trajectory for an additional 500 ms. To obtain target trajectories, neural data were mean-centered and projected onto the top six PCs (computed for that condition). Each target trajectory was normalized by its greatest norm (across times). We trained a total of eighty networks, each with a different weight initialization. The eighty networks included ten each for the two cortical areas, two monkeys, and four condition types.

Network dynamics were governed by:

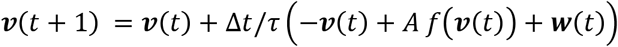

With the learning rule for synaptic input trajectories:

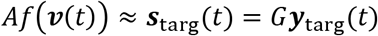

where *f*: = tanh, and 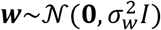 adds noise. ***v*** can be thought of as the membrane voltage and *f*(***v***(*t*)) as the firing rate. *Af*(***v***(*t*)) is then the network input to each unit: the firing rates weighted by the connection strengths. *A* was initialized such that 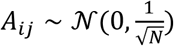 and trained using recursive least squares. ***y***_targ_ is the six-dimensional target trajectory. *G* is an *N* × 6 matrix of random weights, sampled from 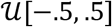, that maps the target trajectory onto a target input of each model unit. The entries of *A* were initialized by draws from a centered normal distribution with variance 1/*N* (where *N* = 50, the number of network units). Simulation employed 4 ms time steps.

To begin a given training epoch, the initial state was set with ***v***(0) based on ***s***_targ_(0) and *A*. The network was simulated, applying recursive least squares^52^ with parameter *α* = 1 to modify *A* as time unfolds. After 1000 training epochs, stability was assessed by simulating the network 100 times, and computing the mean squared difference between the actual and target trajectory. That error was normalized by the variance of the target trajectory, yielding an *R*^2^ value. An average (across the 100 simulated trials) *R*^2^ < 0.9 was considered a failure.

Because population trajectories never perfectly repeated, it was trivially true that networks could follow the full trajectory, for both M1 and SMA, in the complete absence of noise (i.e., for *σ_w_* = 0). For the larger value of *σ_w_* used for our primary analysis, all networks failed to follow the M1 trajectories while most networks successfully followed the SMA trajectories (though there were still some network initializations that never resulted in good solutions). It is of course unclear what value of *σ_w_* is physiologically relevant. We therefore also performed an analysis where we swept the value of *σ_w_* until failure. The level of noise that was tolerated was much greater when networks followed the SMA trajectories. Indeed, some M1 trajectories (for particular conditions) could never be consistently followed even at the lowest noise level tested.

To visualize network activity (Fig 8 b-d) we ‘decoded’ the network population. To do so, we reconstructed the first three dimensions of the trajectory (which should match the first three dimensions of the target trajectory) by inverting *G*.

## Acknowledgements

We thank Y. Pavlova for animal care and Najja J. Marshall who collected a portion of the EMG data. This work was supported by the Grossman Center for the Statistics of Mind, the Simons Foundation (MMC, JPC, LFA), the McKnight Foundation (MMC, JPC), NIH Director’s New Innovator Award DP2 NS083037 (MMC), NIH CRCNS R01NS100066 (MMC and JPC), NIH 1U19NS104649 (MMC, LFA, JPC), NIH 5T32NS064929 (AAR), P30 EY019007, the National Science Foundation (SRB, RK), the Kavli Foundation (MMC), and The Gatsby Charitable Foundation (RK, LFA).

---

**Supplementary Figure 1.**
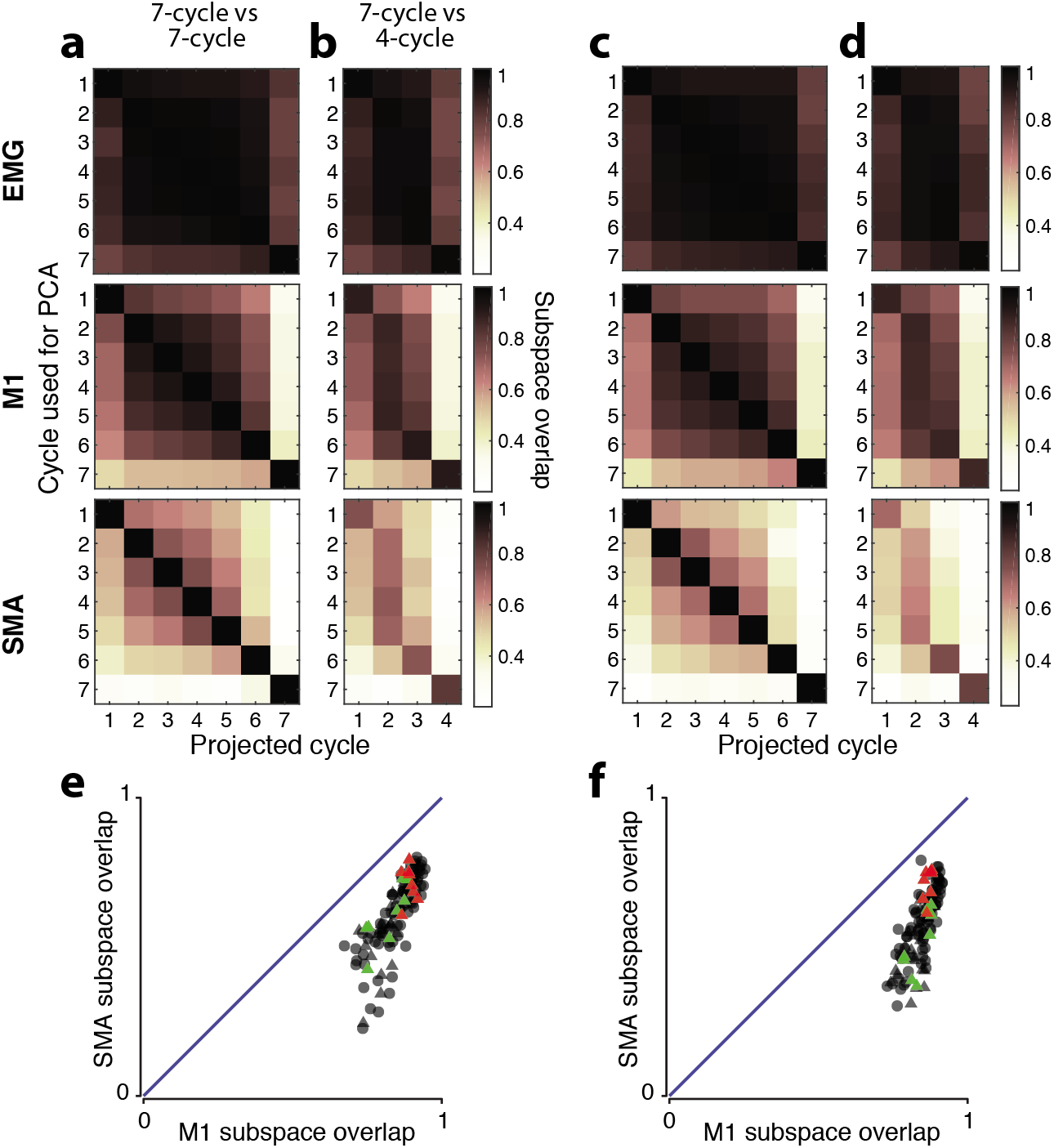
Cycle-to-cycle analysis of subspace overlap in twelve dimensions. Same as Figure 5 but the analysis considered the top twelve PCs. Results are very similar.

**Supplementary Figure 2.**
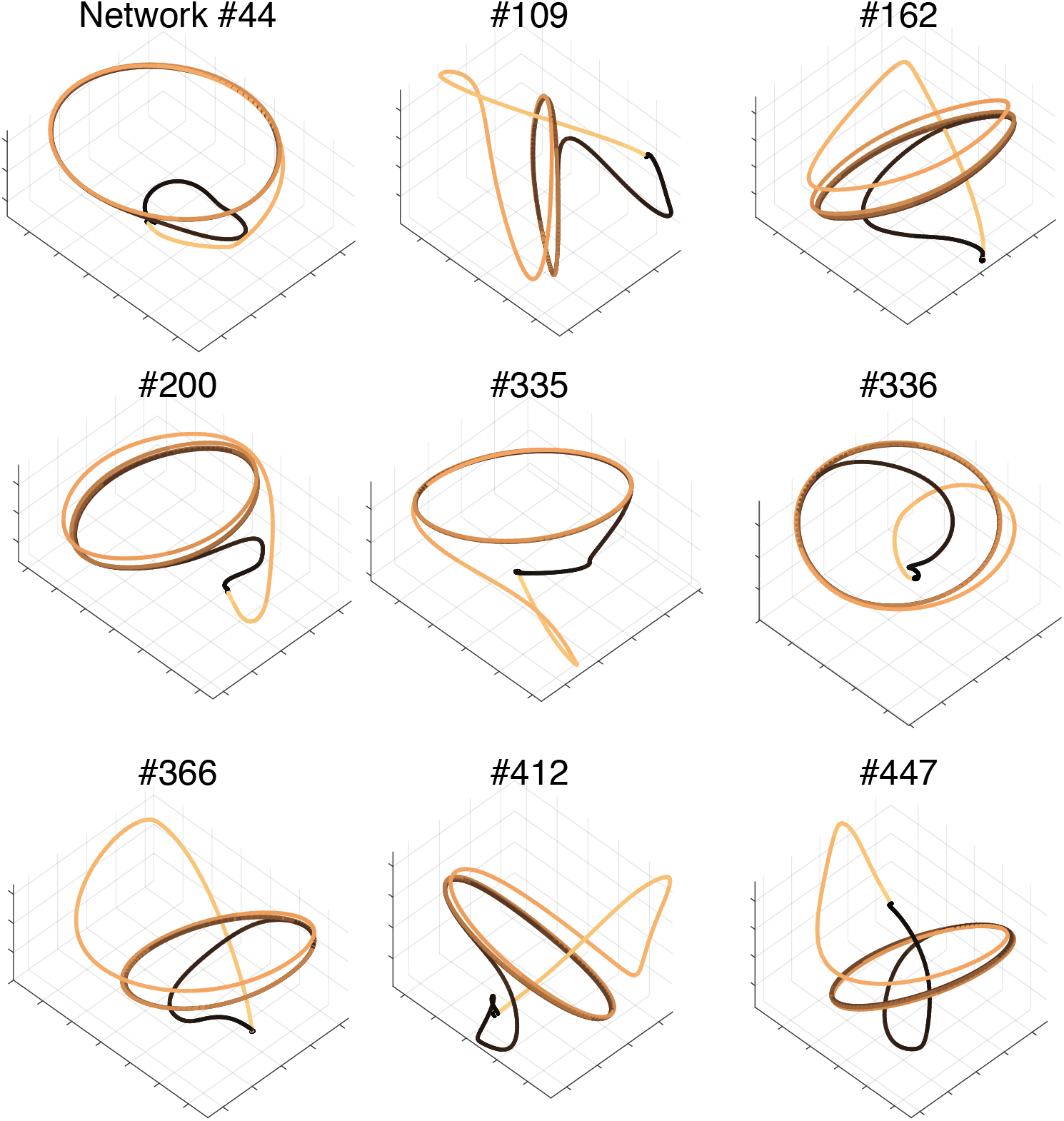
Additional examples of context-naïve network trajectories. Format as for Figure 6a. Nine examples of context-naïve networks trained with different initializations. Activity corresponds to the four-cycle condition.

**Supplementary Figure 3.**
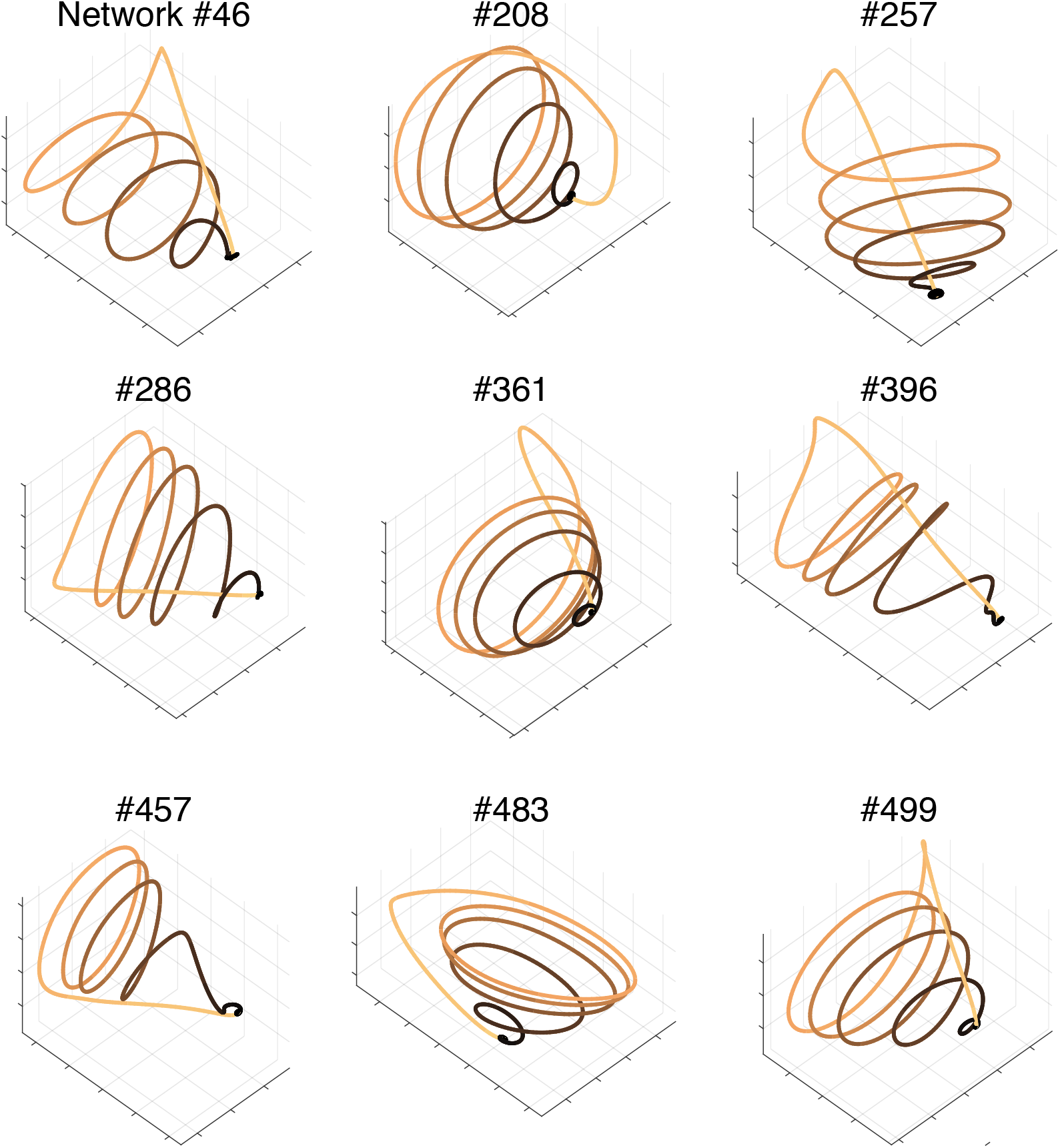
Additional examples of context-tracking. Format as for Figure 6b. Nine examples of context-tracking networks trained with different initializations. Activity corresponds to the four-cycle condition.

**Supplementary Figure 4.**
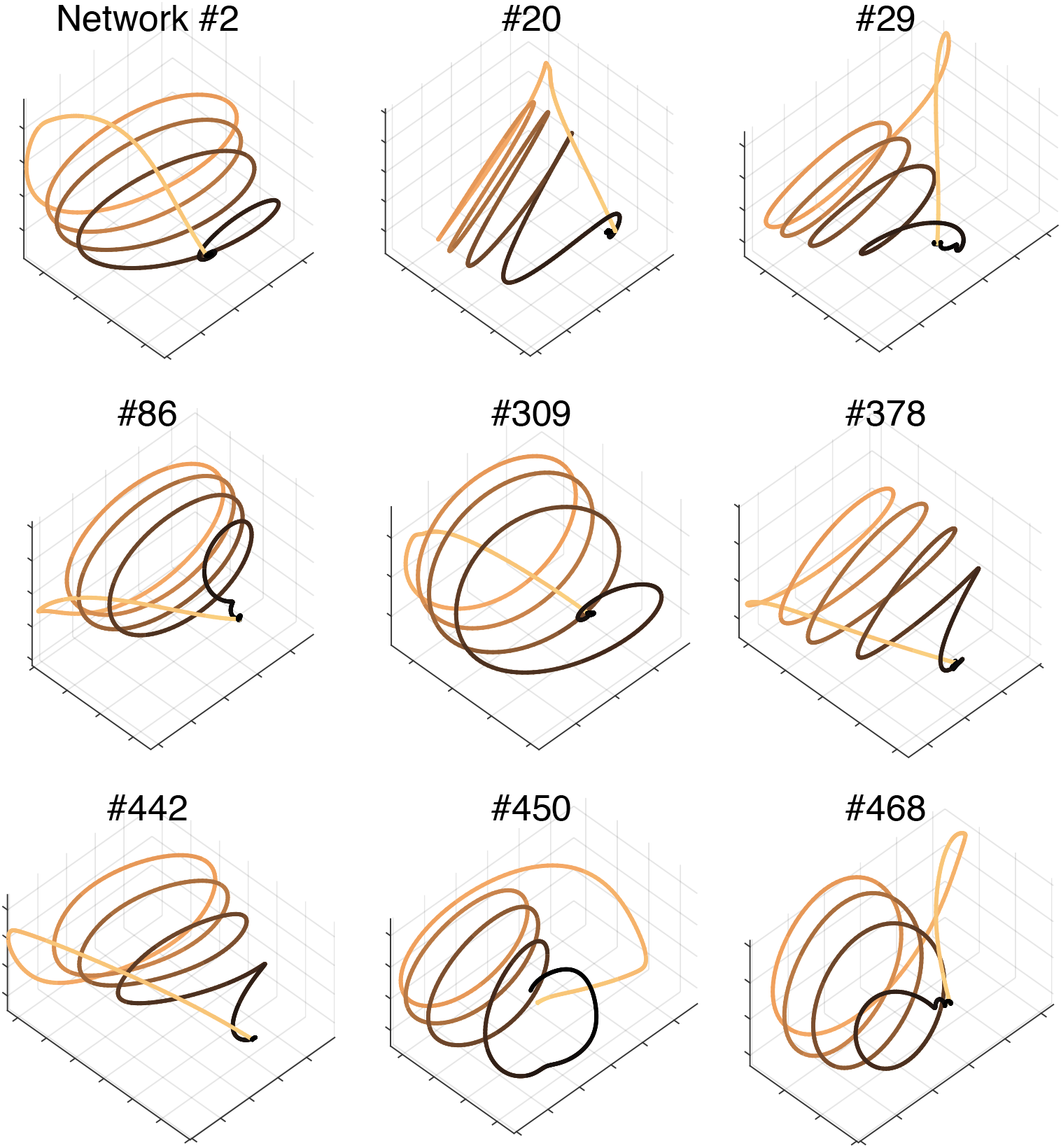
Examples of context-tracking networks trained with a ramping input. Format as for Figure 6b. Nine examples of context-tracking networks trained with different initializations in the presence of a ramping input.

**Supplementary Figure 5.**
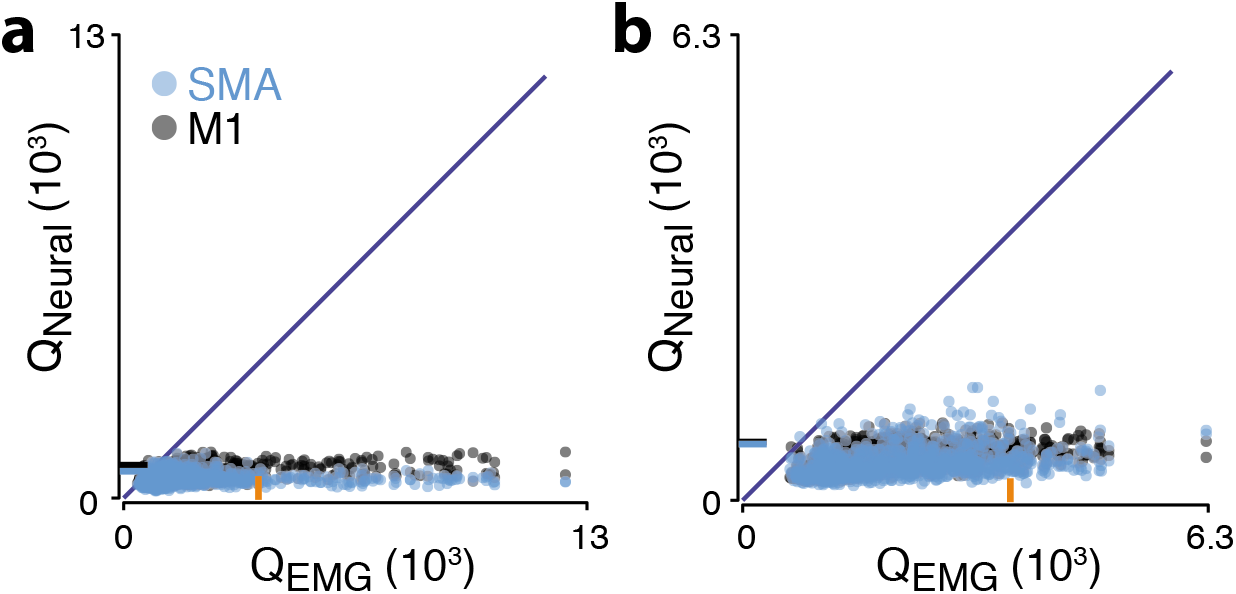
Neural versus muscle population trajectory tangling. a) Neural trajectory tangling versus muscle trajectory tangling (as in Russo et al. 2018) for SMA (blue) and M1 (gray). Both cortical areas show very low trajectory tangling relative to the muscle population trajectories. Data for monkey C. b) Same as panel a but for monkey D.

**Supplementary Figure 6.**
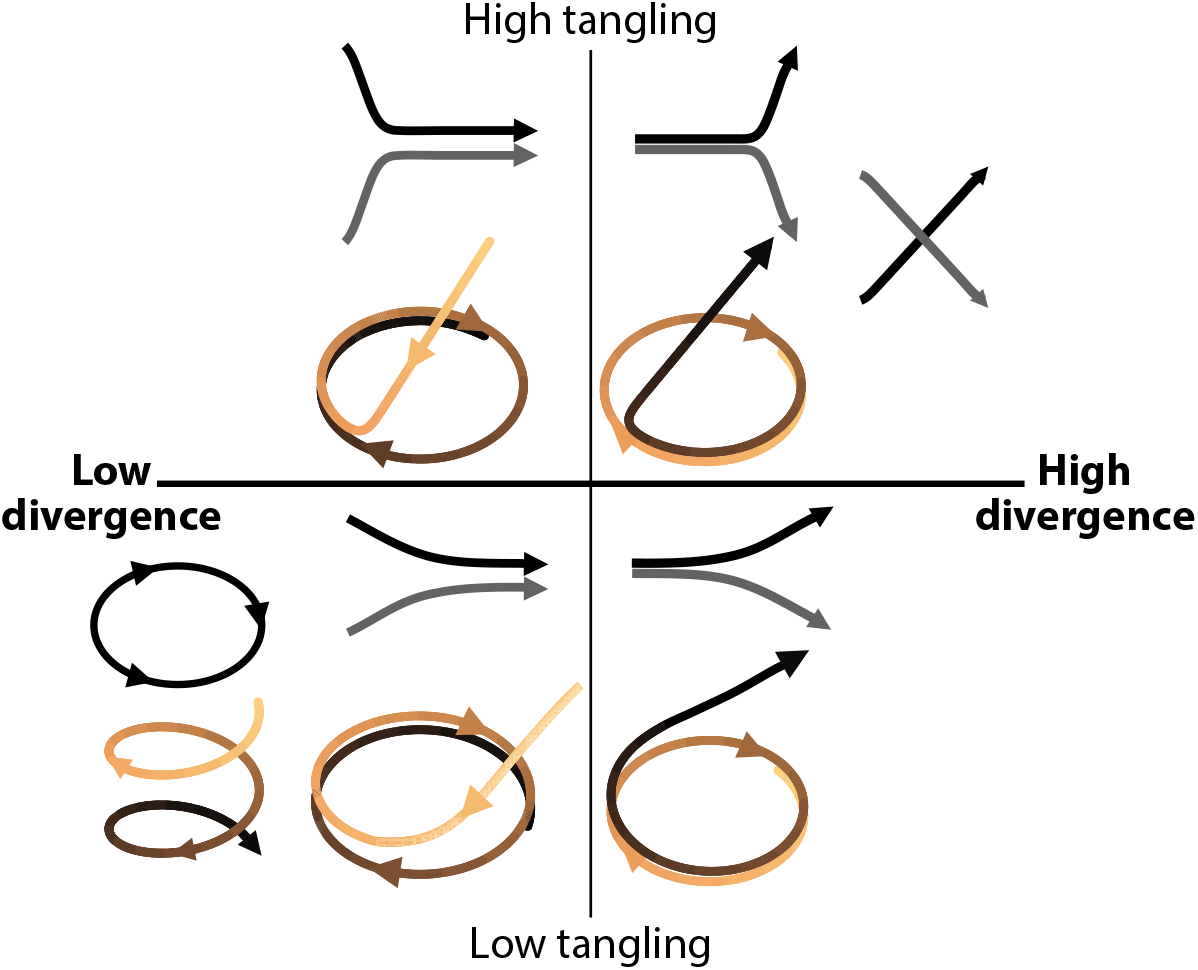
Illustration of trajectories that yield low versus high trajectory divergence and/or trajectory tangling, and consideration of the formal relationship between trajectory divergence and trajectory tanging. Pairs of traces (black and gray) indicate hypothetical trajectories corresponding to different conditions. Single shaded tan-to-black traces indicate hypothetical trajectories for a single condition over time. High trajectory tangling (upper two quadrants) can result from a variety of features, all of which are inconsistent with a smooth underlying flow-field. Not all of these features result in high trajectory divergence. For example, divergence does not become high if trajectories for two conditions converge sharply. High trajectory divergence (right two quadrants) occurs whenever two trajectories (or portions of the same trajectory) are similar at some point in time but later separate. Not all such cases result in high trajectory tangling. For example, tangling remains low if two trajectories diverge slowly. These examples illustrate how trajectory tangling and trajectory divergence assess different aspects of population trajectory geometry. This can also be appreciated formally, by considering their definitions. Trajectory tangling, defined as 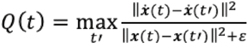, assesses whether the trajectory violates a smooth dynamical flow-field (which would imply similar derivatives for similar states). Trajectory divergence, defined as 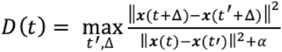, assesses whether two trajectory segments are close but eventually diverge. Unlike tangling, divergence considers future (but not past) events. For example, if trajectories separate slowly, tangling may remain low yet divergence will be high (as in the lower-right quadrant). The relationship between tangling and divergence can be appreciated by considering that the denominators are the same (ignoring constants), and the numerators are related by integration. Consider two quantities. First, ***s***(*τ*) = ***x***(*t* + *τ*) − ***x***(*t*′ + *τ*), the separation between two trajectories at the indicated times. Second, ***v***(*τ*) = ***ẋ***(*t* + *τ*) − ***ẋ***(*t*′ + *τ*), the difference in trajectory velocities. Trajectory divergence is based on 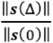. Trajectory tangling is | based on 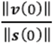. This latter quantity can be modified to consider differences that accumulate over time: 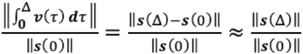 whenever 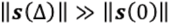. Thus, 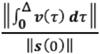 and the divergence metric are nearly identical whenever either is high (differences among small values are irrelevant to our analyses). Thus, divergence can be thought of as a modification of tangling that considers the future, rather than just the present.

**Supplementary Figure 7.**
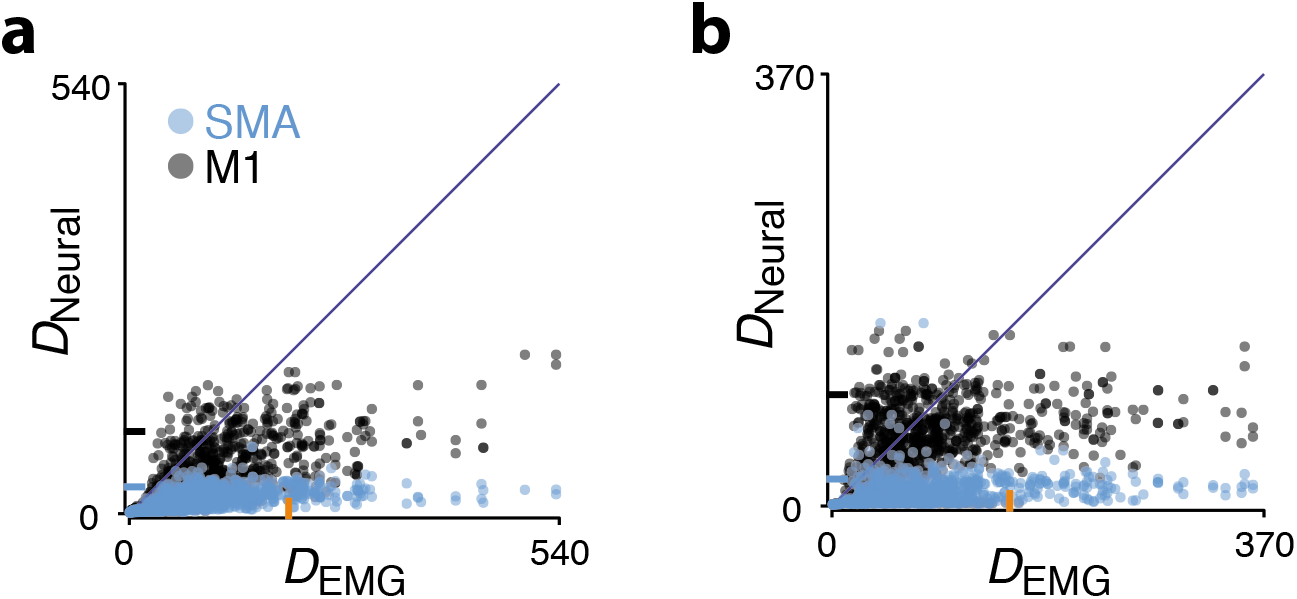
SMA trajectory divergence is low relative to muscle trajectory divergence. **a)** Format as for Figure 7a,b. Blue dots plot SMA trajectory divergence versus trajectory divergence for the muscle population. The orange tick on the horizontal axis plots the 90th percentile of trajectory divergence for the muscle population. The blue tick on the vertical axis plots the same for SMA. SMA trajectory divergence is much lower than muscle trajectory divergence. Black dots plot M1 trajectory divergence versus trajectory divergence for the muscle population. The black tick on the vertical axis plots the 90th percentile of trajectory divergence for M1. This is modestly lower than for the muscles. Data are for monkey C. **b)** Same but for monkey D.

**Supplementary Figure 8.**
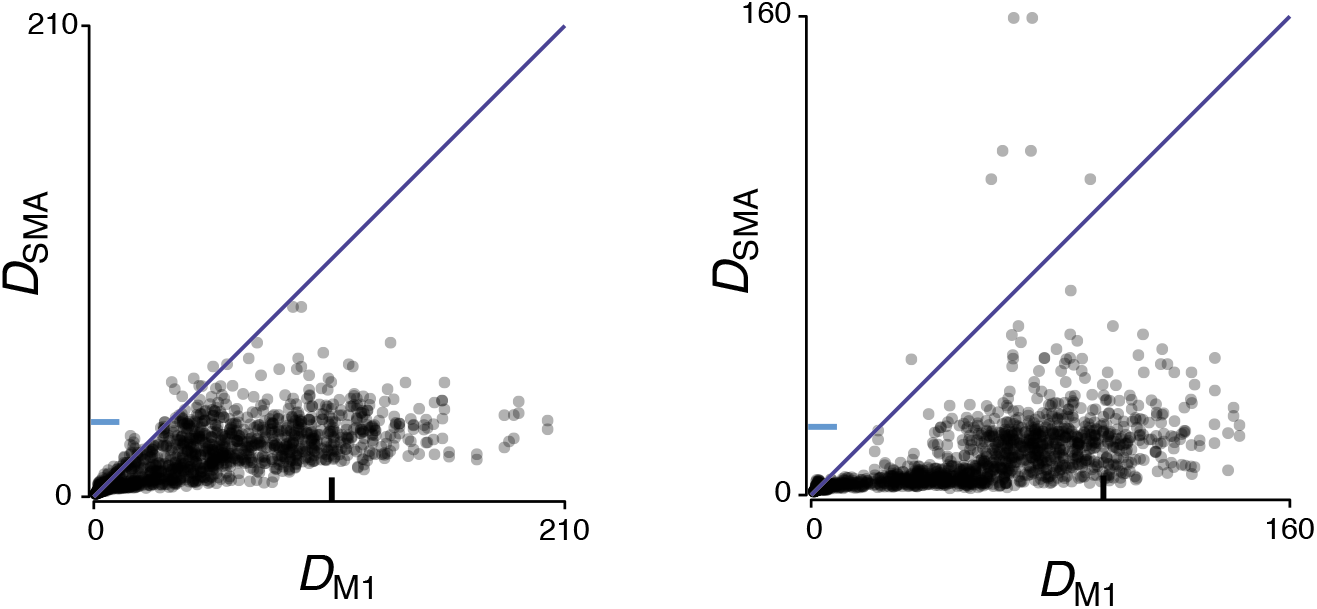
Trajectory divergence in M1 and SMA computed by indexing across all conditions. The analyses of divergence in Figure 7a,b indexed t’ across all times within conditions of the same type: e.g., across the four distances when cycling forward starting from the top. Those analyses thus assessed whether SMA trajectories track contextual factors related to the number of cycles to be performed. If SMA does so, divergence will be low. One can also ask whether trajectory divergence remains low when considered across all conditions. For example, is SMA activity during steady-state cycles different depending on whether the movement started at the bottom (and will thus end at the bottom) or started at the top (and will thus end at the top). Steady state cycles for bottom-start and top-start conditions have essentially identical patterns of muscle activity, but future muscle activity (on the last cycle and after stopping) will be different. To ask whether divergence remains low when considering all conditions (including both cycling directions) together, the present analysis indexes t’ across all times within all conditions. Same plotting format as Figure 7a,b. Left and right panels plot data for monkeys C and D.

## References

1 Roland, P. E., Larsen, B., Lassen, N. A. & Skinhoj, E. Supplementary Motor Area and Other Cortical Areas in Organization of Voluntary Movements in Man. Journal of Neurophysiology 43, 118–136 (1980).

2 Penfield, W. & Welch, K. The Supplementary Motor Area of the Cerebral Cortex - a Clinical and Experimental Study. Ama Arch Neurol Psy 66, 289–317, doi:DOI 10.1001/archneurpsyc.1951.02320090038004 (1951).

3 Eccles, J. C. The Initiation of Voluntary Movements by the Supplementary Motor Area. Arch Psychiat Nerven 231, 423–441, doi:Doi 10.1007/Bf00342722 (1982).

4 Laplane, D., Talairach, J., Meininger, V., Bancaud, J. & Orgogozo, J. M. Clinical Consequences of Corticectomies Involving Supplementary Motor Area in Man. J Neurol Sci 34, 301–314, doi:Doi 10.1016/0022-510x(77)90148-4 (1977).

5 Krainik, A. et al. Role of the supplementary motor area in motor deficit following medial frontal lobe surgery. Neurology 57, 871–878 (2001).

6 Boccardi, E., Della Sala, S., Motto, C. & Spinnler, H. Utilisation behaviour consequent to bilateral SMA softening. Cortex 38, 289–308 (2002).

7 Nakamura, K., Sakai, K. & Hikosaka, O. Neuronal activity in medial frontal cortex during learning of sequential procedures. Journal of Neurophysiology 80, 2671–2687 (1998).

8 Shima, K. & Tanji, J. Both supplementary and presupplementary motor areas are crucial for the temporal organization of multiple movements. J Neurophysiol 80, 3247–3260, doi:10.1152/jn.1998.80.6.3247 (1998).

9 Mushiake, H., Inase, M. & Tanji, J. Neuronal activity in the primate premotor, supplementary, and precentral motor cortex during visually guided and internally determined sequential movements. J Neurophysiol 66, 705–718, doi:10.1152/jn.1991.66.3.705 (1991).

10 Tanji, J. & Mushiake, H. Comparison of neuronal activity in the supplementary motor area and primary motor cortex. Brain Res Cogn Brain Res 3, 143–150 (1996).

11 Tanji, J. & Kurata, K. Comparison of movement-related activity in two cortical motor areas of primates. J Neurophysiol 48, 633–653, doi:10.1152/jn.1982.48.3.633 (1982).

12 Boudrias, M. H., Belhaj-Saif, A., Park, M. C. & Cheney, P. D. Contrasting properties of motor output from the supplementary motor area and primary motor cortex in rhesus macaques. Cereb Cortex 16, 632–638, doi:10.1093/cercor/bhj009 (2006).

13 Sohn, J. W. & Lee, D. Order-dependent modulation of directional signals in the supplementary and presupplementary motor areas. J Neurosci 27, 13655–13666, doi:10.1523/JNEUROSCI.2982-07.2007 (2007).

14 Thaler, D., Chen, Y. C., Nixon, P. D., Stern, C. E. & Passingham, R. E. The functions of the medial premotor cortex. I. Simple learned movements. Exp Brain Res 102, 445–460 (1995).

15 Remington, E. D., Narain, D., Hosseini, E. A. & Jazayeri, M. Flexible Sensorimotor Computations through Rapid Reconfiguration of Cortical Dynamics. Neuron 98, 1005-+, doi:10.1016/j.neuron.2018.05.020 (2018).

16 Wang, J., Narain, D., Hosseini, E. A. & Jazayeri, M. Flexible timing by temporal scaling of cortical responses. Nat Neurosci 21, 102-+, doi:10.1038/s41593-017-0028-6 (2018).

17 Tanji, J. & Shima, K. Role for supplementary motor area cells in planning several movements ahead. Nature 371, 413–416, doi:10.1038/371413a0 (1994).

18 Kornysheva, K. & Diedrichsen, J. Human premotor areas parse sequences into their spatial and temporal features. Elife 3, e03043, doi:10.7554/eLife.03043 (2014).

19 Shima, K. & Tanji, J. Neuronal activity in the supplementary and presupplementary motor areas for temporal organization of multiple movements. J Neurophysiol 84, 2148–2160, doi:10.1152/jn.2000.84.4.2148 (2000).

20 Cadena-Valencia, J., Garcia-Garibay, O., Merchant, H., Jazayeri, M. & de Lafuente, V. Entrainment and maintenance of an internal metronome in supplementary motor area. Elife 7, doi:10.7554/eLife.38983 (2018).

21 Gamez, J., Mendoza, G., Prado, L., Betancourt, A. & Merchant, H. The amplitude in periodic neural state trajectories underlies the tempo of rhythmic tapping. PLoS Biol 17, e3000054, doi:10.1371/journal.pbio.3000054 (2019).

22 Saxena, S. & Cunningham, J. P. Towards the neural population doctrine. Current opinion in neurobiology 55, 103–111, doi:10.1016/j.conb.2019.02.002 (2019).

23 Gallego, J. A. et al. Cortical population activity within a preserved neural manifold underlies multiple motor behaviors. Nat Commun 9, 4233, doi:10.1038/s41467-018-06560-z (2018).

24 Pandarinath, C. et al. Latent Factors and Dynamics in Motor Cortex and Their Application to Brain-Machine Interfaces. J Neurosci 38, 9390–9401, doi:10.1523/JNEUROSCI.1669-18.2018 (2018).

25 Morrow, M. M. & Miller, L. E. Prediction of muscle activity by populations of sequentially recorded primary motor cortex neurons. J Neurophysiol 89, 2279–2288, doi:10.1152/jn.00632.2002 (2003).

26 Sussillo, D., Churchland, M. M., Kaufman, M. T. & Shenoy, K. V. A neural network that finds a naturalistic solution for the production of muscle activity. Nat Neurosci 18, 1025-+, doi:10.1038/nn.4042 (2015).

27 DiCarlo, J. J., Zoccolan, D. & Rust, N. C. How does the brain solve visual object recognition? Neuron 73, 415–434, doi:10.1016/j.neuron.2012.01.010 (2012).

28 Schaffelhofer, S. & Scherberger, H. Object vision to hand action in macaque parietal, premotor, and motor cortices. Elife 5, doi:10.7554/eLife.15278 (2016).

29 Foster, J. D. et al. A freely-moving monkey treadmill model. J Neural Eng 11, 046020, doi:10.1088/1741-2560/11/4/046020 (2014).

30 Stopfer, M. & Laurent, G. Short-term memory in olfactory network dynamics. Nature 402, 664–668, doi:Doi 10.1038/45244 (1999).

31 Remington, E. D., Egger, S. W., Narain, D., Wang, J. & Jazayeri, M. A Dynamical Systems Perspective on Flexible Motor Timing. Trends Cogn Sci 22, 938–952, doi:10.1016/j.tics.2018.07.010 (2018).

32 Sussillo, D. & Barak, O. Opening the Black Box: Low-Dimensional Dynamics in High-Dimensional Recurrent Neural Networks. Neural Comput 25, 626–649, doi:Doi 10.1162/NECO_a_00409 (2013).

33 Raposo, D., Kaufman, M. T. & Churchland, A. K. A category-free neural population supports evolving demands during decision-making. Nat Neurosci 17, 1784–1792, doi:10.1038/nn.3865 (2014).

34 Hall, T. M., de Carvalho, F. & Jackson, A. A common structure underlies low-frequency cortical dynamics in movement, sleep, and sedation. Neuron 83, 1185–1199, doi:10.1016/j.neuron.2014.07.022 (2014).

35 Michaels, J. A., Dann, B. & Scherberger, H. Neural Population Dynamics during Reaching Are Better Explained by a Dynamical System than Representational Tuning. PLoS Comput Biol 12, e1005175, doi:10.1371/journal.pcbi.1005175 (2016).

36 Ames, K. C., Ryu, S. I. & Shenoy, K. V. Neural Dynamics of Reaching following Incorrect or Absent Motor Preparation. Neuron 81, 438–451, doi:10.1016/j.neuron.2013.11.003 (2014).

37 Driscoll, L. N., Golub, M. D. & Sussillo, D. Computation through Cortical Dynamics. Neuron 98, 873–875, doi:10.1016/j.neuron.2018.05.029 (2018).

38 Russo, A. A. et al. Motor Cortex Embeds Muscle-like Commands in an Untangled Population Response. Neuron 97, 953-+, doi:10.1016/j.neuron.2018.01.004 (2018).

39 Merchant, H. & de Lafuente, V. Introduction to the neurobiology of interval timing. Adv Exp Med Biol 829, 1–13, doi:10.1007/978-1-4939-1782-2_1 (2014).

40 Romo, R. & Schultz, W. Role of primate basal ganglia and frontal cortex in the internal generation of movements. III. Neuronal activity in the supplementary motor area. Exp Brain Res 91, 396–407 (1992).

41 Nachev, P., Kennard, C. & Husain, M. Functional role of the supplementary and presupplementary motor areas. Nat Rev Neurosci 9, 856–869, doi:10.1038/nrn2478 (2008).

42 Lara, A. H., Cunningham, J. P. & Churchland, M. M. Different population dynamics in the supplementary motor area and motor cortex during reaching. Nat Commun 9, 2754, doi:10.1038/s41467-018-05146-z (2018).

43 Hatsopoulos, N. G., Paninski, L. & Donoghue, J. P. Sequential movement representations based on correlated neuronal activity. Exp Brain Res 149, 478–486, doi:10.1007/s00221-003-1385-9 (2003).

44 Yokoi, A., Arbuckle, S. A. & Diedrichsen, J. The Role of Human Primary Motor Cortex in the Production of Skilled Finger Sequences. J Neurosci 38, 1430–1442, doi:10.1523/JNEUROSCI.2798-17.2017 (2018).

45 Mante, V., Sussillo, D., Shenoy, K. V. & Newsome, W. T. Context-dependent computation by recurrent dynamics in prefrontal cortex. Nature 503, 78-+, doi:10.1038/nature12742 (2013).

46 Churchland, M. M. et al. Neural population dynamics during reaching. Nature 487, 51-+, doi:10.1038/nature11129 (2012).

47 Kaufman, M. T., Churchland, M. M., Ryu, S. I. & Shenoy, K. V. Cortical activity in the null space: permitting preparation without movement. Nat Neurosci 17, 440–448, doi:10.1038/nn.3643 (2014).

48 Gallego, J. A., Perich, M. G., Miller, L. E. & Solla, S. A. Neural Manifolds for the Control of Movement. Neuron 94, 978–984, doi:10.1016/j.neuron.2017.05.025 (2017).

49 Seely, J. S. et al. Tensor Analysis Reveals Distinct Population Structure that Parallels the Different Computational Roles of Areas M1 and V1. PLoS Comput Biol 12, e1005164, doi:10.1371/journal.pcbi.1005164 (2016).

50 Werbos, P. J. Generalization of backpropagation with application to a recurrent gas market model. Neural Networks 1, 339–356 (1988).

51 DePasquale, B., Cueva, C. J., Rajan, K., Escola, G. S. & Abbott, L. F. full-FORCE: A target-based method for training recurrent networks. PLoS One 13, e0191527, doi:10.1371/journal.pone.0191527 (2018).

52 Sussillo, D. & Abbott, L. F. Generating coherent patterns of activity from chaotic neural networks. Neuron 63, 544–557 (2009).

